# The Impact of Charge Regulation and Ionic Intranuclear Environment on the Nucleosome Core Particle

**DOI:** 10.1101/2024.11.11.623012

**Authors:** Rikkert J. Nap, Paola Carrillo Gonzalez, Aria E. Coraor, Ranya K. A. Virk, Juan de Pablo, Vadim Backman, Igal Szleifer

## Abstract

We theoretically investigate how the intranuclear environment influences the charge of a nucleosome core particle (NCP) - the fundamental unit of chromatin consisting of DNA wrapped around a core of histone proteins. The molecular-based theory explicitly considers the size, shape, conformations, charges, and chemical states of all molecular species - thereby linking the structural state with the chemical/charged state of the system. We investigate how variations in monovalent and divalent salt concentrations, as well as pH, affect the charge distribution across different regions of an NCP and quantify the impact of charge regulation. The effective charge of an NCP emerges from a delicate and complex balance involving the chemical dissociation equilibrium of the amino acids and the DNA-phosphates, the electrostatic interaction between them, and the translational entropy of the mobile solution ions, i.e., counter ion release and ion condensation. From our results, we note the significant effect of divalent magnesium ions on the charge and electrostatic energy as well as the counterion cloud that surrounds an NCP, as a function of magnesium concentration, charge neutralization, and even charge inversion is predicted - in line with experimental observation of NCPs. The strong Mg-dependence of the nucleosome charge state arises from ion bridges between two DNA-phosphates and one Mg ^+^ ion. We demonstrate that to describe and predict the charged state of an NCP properly, it is essential to consider molecular details, such as DNA-phosphate ion condensation and the acid-base equilibrium of the amino acids that comprise the core histone proteins.

## I. INTRODUCTION

Electrostatic interactions in their many forms play a fundamental role across different nanoscopic^1^and biological systems^2^. Chromatin, the supra-molecular complex composed of DNA polyacids and histone proteins found in the nucleus, carries the genetic code that dictates proper cell function and is a prime example of a biological system where electrostatic interactions are important^3^. DNA carries a negative charge due to the sugar-phosphate backbone, while the chargeable amino acids of the histone proteins imbue the core protein with a net positive charge. However, the amino acid residues of histones only partially neutralize the highly negative charge of DNA. Hence, chromatin (usually) carries an overall net negative charge. Consequently, the electrostatic interactions between the chargeable amino acids and the phosphate backbone charges affect the stability and packing of chromatin. As structure dictates DNA accessibility to the transcription machinery, gene expression, and cell function assembly are consequently influenced by charges as well.

Numerous experiments and simulations have shown that chromatin organization is influenced by the aggregation and phase separation of chromatin, which are driven by self-interactions. Considering the polyelectrolyte characteristics of chromatin, these self-interactions are also significantly modulated by monovalent and divalent salt concentrations. These influences range from local properties, such as persistence length, to the aggregation and phase separation of chromatin, as well as the compaction of chromatin domains, and even to cell fate^4–11^. For example, in vitro experiments have highlighted the aggregation of nucleosome core particles (NCPs) as a function of pH, ionic concentrations, and ionic valency using techniques like turbidity and UV-adsorption measurements^12–15^. Additionally, the compaction of reconstituted nucleosomes with increasing levels of divalent ions has been directly visualized using fluorescence microscopy^7,16^. In vivo heat shock experiments have also noted intercellular acidification (pH changes) and changes in transcription patterns of cells^17^, indicating that ions play a major role in chromatin remodeling. For a comprehensive review of past chromatin aggregation and phase-separation studies, the reader is referred to a recent review by Hansen et al.^11^. Despite an understanding of the basic physics involved, significant questions remain regarding the interplay between electrostatic interactions and chromatin structure. Specifically, how is chromatin organized inside the nucleus while mitigating the electrostatic repulsions caused by the charges in the system? How is the intranuclear environment—including ions, pH, and molecular crowding such as chromatin density— implicated in the regulation of chromatin structure and charge?

Chromatin conformation plays a crucial role in the regulation of gene expression and has been considered as a therapeutic target^18^ and chromatin structure is expected to depend in part on ionic interactions. Thus, understanding how ions and pH mechanistically regulate chromatin structure is fundamentally important and has potential therapeutic applications.

Abnormal chromatin organization underlies a wide spectrum of conditions, including cancer, neurological disorders, and developmental abnormalities. Leveraging how ions influence chromatin structure and function, researchers could identify specific molecular targets for therapeutic intervention. Maeshima et al., for example, observed that ATP inhibits Mg^2+^-dependent chromatin condensation in vitro, suggesting a physical pathway by which enzymes and kinases could indirectly affect ion concentration and electrostatic interactions, and thus molding chromatin structure and influencing gene expression. Here, we envision that understanding the fundamental mechanisms behind ions and their role in chromatin organization presents a promising avenue for the development of novel therapeutic strategies targeting chromatin-associated diseases and disorders.

Despite these insights, a complete understanding of the principal mechanisms of chromatin charging and packing remains elusive. This is partly due to the technical limitations of current experiments, which cannot concomitantly measure all relevant properties of the system. To address these challenges, researchers turn to computational approaches, which can model and simulate complex interactions and provide a more comprehensive view of chromatin dynamics. Numerous computational approaches, ranging from all-atomistic to coarse-grained simulations, have helped to uncover the effects of ions on chromatin mechanistically^19^. Still, computational approaches also have challenges and limitations. While the role of charges on the amino acid and phosphates has been taken into account to various degrees^20–36^, there have been few efforts to theoretically model how acid-base reactions, ion condensation, and divalent ion bridging as well as solution factors like pH and salt concentrations, affect the organization of chromatin.

Furthermore, performing atomistic or MD simulations to describe chromatin is computationally challenging due to the vast number of atoms involved. For example, a canonical nucleosome consists of approximately *O*(10^4^) atoms, making full atomistic simulation CPU costly. While atomistic and mesoscale coarse-grained MD simulations provide many molecular details, they generally impose implicit ions and solvent, a fixed charge distribution, and do not consider the possibility of dynamic chemical equilibria between protonation and deprotonation of acids and bases nor the explicit possibility of ion condensation or ion bridging. Acid-base chemical equilibrium can be introduced in MD simulations using constant-pH or reaction ensemble simulations^37,38^, but this is difficult and costly, in development- and CPU-time, to implement except in small, relatively simple systems^37,39^.

Another limitation is that MD simulations, due to their time-consuming nature, are not practical for systematic variation of parameters like salt concentration and pH. Additionally, simulations involving multivalent ions are hampered by the occurrence of long-lived metastable kinetically trapped states and ion pairs which can hinder accurate modeling.^40^ Incorporating Mg^2+^ interactions in implicit coarse-grained simulation schemes is challenging.^4^. In summary, most computations are often performed for fixed charges, implicit ions, and implicit solvent, fixed pH, meaning that proton and ion concentrations are position-independent, and employ a coarse-grained representation of the nucleosome. Notable exceptions are recent investigations by Lin et al and Sun et al who considered the effect of explicit monovalent and divalent (Mg^2+^) ions respectively using coarse-grained simulations for nucleosome arrays with fixed histone charges^28,41,42^.

Therefore, we introduce a detailed Molecular Theory (MT) approach to investigate how the intranuclear environment influences the charge of an NCP with and without histone tails. MT explicitly considers the size, shape, conformations, charges, and chemical states of all molecular species, thus linking the structural state with the chemical or charged state of the system. Importantly, using MT, it is computationally feasible to cover a wide range of environmental conditions. Here, we use the MT approach to determine how the charge and electrostatic interaction of a single nucleosome change in response to varied environmental conditions. We examined how the effective charge of an NCP depends on monovalent and divalent salt concentrations and pH.

Since the effects of acid-base reaction equilibrium, varied pH, ion condensation, and the ionic strength of the electrolyte solution on an NCP remains largely unexplored, it is our goal in this study to elucidate the details of this dependence and provide deeper insight into the ionic regulation of an NCP in a realistic nuclear environment.

We explicitly include the chemical equilibrium between the protonated, deprotonated, and ion-condensed states of the chargeable DNA-phosphates and amino acid (AA) comprising the histone core protein. Importantly, the theory does not assume the charged state of the DNA and the AA residues of the nucleosome but rather predicts the position-dependent state of a charge. The theory is based on a molecular statistical thermodynamic approach that has previously been developed to predict thermodynamic and structural properties of end-tethered polymers and weakly ionizable polyelectrolyte, see review^43^ and references therein. Predictions of the MT approach have been found to agree with experimental observations for relevant biological systems, including polyelectrolyte brushes, nuclear pore complexes, and phenomena such as protein adsorption on polymer brush surfaces^44,45^ The theory was also able to predict the charges of the biopolymer aggrecan, bacteriophages, and ligated nanoparticles^46–48^. Importantly, this theoretical approach does not assume a predefined nucleosome charge, such as by ideal solution behavior. Instead, the nucleosome’s charge distribution emerges from the free energy minimization.

Herein, we would like to address the effects of charge regulation on the charge and electrostatic interaction of an NCP with and without histone tails to gain a fundamental understanding of how the intranuclear physicochemical environment of monovalent and divalent ions modulate the charge. This will be a first step toward a more proper fundamental understanding of the effect of charge regulation of chromatin in the nucleus.

## II. MODEL

Here, we present a molecular theoretical approach for describing a single nucleosome with and without histone tails in an aqueous solution. We consider a nucleosome interacting with a nuclear environment defined by a given pH and containing monovalent KCl and NaCl as well as divalent MgCl_2_ ions at physiological concentrations. The ions in the system are assumed to be completely dissociated. Although the nucleus contains additional cations, such as polyamines, these salts were selected because they are the most common intracellular ions. We select and vary both pH and salt concentrations around physiological relevant ranges found in experiments. In our calculations, the pH of the reservoir is adjusted by adding either HCl or NaOH^49^.

### A. Molecular Representation of the Nucleosome

In the NCP, is represented by a triplet of coarse-grained units, namely a chargeable phosphate (P), a sugar (S), and a nucleobase, either (C, T, A, or G) using the 3SPN.2 model developed by the de Pablo group^50^. The histone core (octamer) proteins, composed of AAs, are represented at the atomic level. Each atom in the side chain of an AA is depicted as a single unit. Assuming the histone protein is in a frozen ‘crystal’ state we can use the atom positions determined from X-ray diffraction experiments directly in our model^51^. Many coarse-grained models represent a single amino acid with one effective unit/sphere. In doing so usually, the alpha carbon is selected as both volume and charge center. However in large amino acids, the alpha carbon location and charge center do not coincide. For example, the alpha carbon and the charge center are roughly 0.6 *nm* apart in lysine, one of the more prevalent AAs of the histone octamer. In including a detailed representation of the core histone, we can account more accurately for the charge localization as well as the asymmetric shape and excluded volume interactions of the amino acids.

### B. Treatment of ions and complexation

The charge of the nucleosome arises from a plethora of chargeable moieties present, which include the chargeable amino residues of the core histone proteins and the phosphates of the wrapped DNA chain. Here deprotonation of the carboxylate group

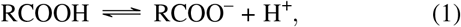

of the side chains in aspartic and glutamic acid and deprotonation of the hydroxyl in the phenyl group of tyrosine

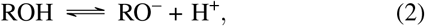

contribute negative charges to the histone. Similarly, protonation of the amine groups of the side chains of arginine, histidine, and lysine results in positive charges.

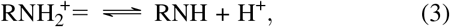

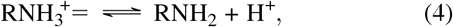

Cysteine has a thiol functional end group that is a weakly acidic,

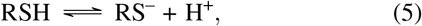

however, cysteine is usually not considered to be an acid since the thiol group is often reactive and can form disulfide bonds. Theoretically, we can consider two different cases in which either the acid-base equilibrium of cysteine was considered or ignored. Compared to other more abundant chargeable amino acids, there are only two cysteines in the entire structure. Consequently, the influence of cysteine on the total effective charge of a nucleosome will be small irrespective of whether the acid-base equilibrium is considered or not.

Besides acid-base equilibrium, we also explicitly include the process of ion-pairing or ion-condensation of oppositely charged ions with acidic AAs

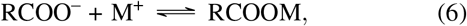

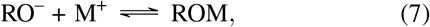

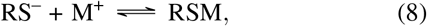

Here *M*^+^ represents the monovalent cations in the system, *Na*^+^ or *K*^+^ that can bind to negatively charged acids. Similarly, a positively charged amino-acid base (amine and amide) can form an ion-pair with *Cl*^*−*^

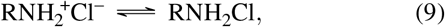

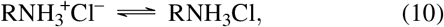

The chemical reactions above represent physical interactions associated with short-range electrostatic interactions originating from ion-binding and ion-pairing. In representing ion binding as a chemical reaction, we provide a convenient and transparent way of including short-range electrostatic interaction in the theory. Explicitly incorporating ion pairing provides a clearer understanding of the charge regulation process that occurs within nucleosome systems.

Like the chargeable AAs, the phosphate groups of the DNA base pairs form a source of negative charge and contribute to the total effective charge of the nucleosome. The DNA-phosphate is assumed to be in one of six different chemical states: deprotonated (P^−^), protonated (PH) or condensed with K^+^, Na^+^, or Mg^2+^ counter ions. These different chemically charged states occur through the following chemical reactions that are explicitly included in the theory

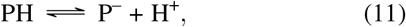

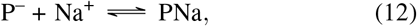

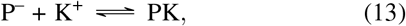

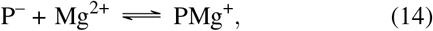

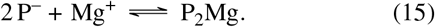

The condensed states of monovalent cations are denoted as PNa, PK. Given its divalent characteristics, there are two different condensed states for Mg^2+^: 1:1 binding, PMg^+^, and 2:1 binding, P_2_Mg, of the phosphates with the divalent cations. The last reaction, in which two phosphates are bound with one magnesium or ion bridging reaction, does not occur directly but via the reaction P^−^ + PMg^+^ ⇌ P_2_Mg. However, these reactions are thermodynamically equivalent. We have not considered the formation of ion pairs involving multiple divalent cations and phosphates simultaneously. Thus, monovalent cations can only bind with one phosphate, while divalent cations can form additional ion bridges with a stoichiometry of two phosphates to one Mg^2+^ ion.

In an ideal solution, the activity coefficients of the products and reactants are one, and the extent of each chemical reaction is determined solely by Δ*G*^⦵^), the standard Gibbs free energy of each reaction or equivalently the chemical equilibrium constants (*K*^⦵^) as well as the concentration of the involved moieties, namely the concentration of AAs and phosphates and the reservoir concentration of protons and ions. Ideal solutions imply that the moiety involved in the chemical reaction are in infinite dilution and not interacting with each other. However, the chargeable AAs and phosphates are not in ideal conditions. They are part of a closely packed nucleosome, experiencing large osmotic and electrostatic interactions. This will cause the charging of the AAs and phosphates to deviate considerably from ideal solution behavior. In the next section, we derive and present the non-ideal chemical reaction equations that occur for the AAs and the DNA-phosphates.

## C. Molecular Theory Free Energy

The MT free energy describes a single nucleosome in contact with an aqueous electrolyte solution and has several distinct contributions which can be summarized as follows

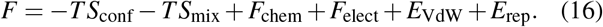

The first contribution (*S*_conf_) is related to the conformational entropy of the nucleosome. The second term encompasses the mixing or translational entropy of the mobile ions and solvent (*S*_mix_). The next two contributions stem from the acid-base chemical equilibrium and counterion condensation of the amino acid and the phosphates (*F*_chem_) and the electrostatic energy (*F*_elect_). The subsequent term(*F*_VdW_) represents the effective Van der Waals, or hydrophobic interactions, among the DNA and amino acid residues. The last term (*E*_rep_) accounts for the steric repulsions, or excluded volume interactions, among all molecular species.

Here, we focus on the most important feature of the free energy of the current study: namely the free energy contribution related to the acid-base equilibria and ion condensations. The remaining terms, such as the translational entropy of solvent and mobile ions as well as the electrostatic energy have been discussed previously in the context of end-tethered polyelectrolytes. For a detailed discussion of these terms, the reader is referred to the supporting materials and previous publications.^43,52^ The supporting material presents the complete free energy functional.

Excluded volume interactions between molecules are incorporated by assuming the system is incompressible at every point. As a result, the sum of the volume fractions of all molecular species is equal to one at each position.

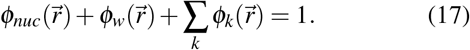

These packing constraints are enforced by introducing Lagrange multipliers 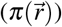. Here 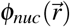, corresponds to the total volume fraction of the nucleosome while the second and third term relates to the volume fraction of the (water) solvent and all mobile ionic species. The latter two volume fractions are given by their number density times their volume: 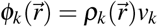. Here, we assume that the single nucleosome is a frozen “state” for both the core AAs and the DNA. As a result, the conformational entropy *S*_*con f*_ = 0 and the number density distribution of all molecular species comprising the nucleosome, 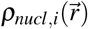, are fixed. In maintaining the core AAs and DNA-phosphates in a single state, we focus on the charge regulation effects of a tail-less NCP. This approach excludes the consideration of multiple histone octamers and the influence of disordered tails as additional conformations would be necessary. It is worth noting that atom positions determined from the X-ray diffraction experiments^51^ included one set of positions of the disordered histone tails. To explore the effect of disordered tails, we considered the NCP with the disordered tails intact, using the experimentally reported positions. However, this should be regarded as an additional approximation since the disordered histone tails, by definition, may vary positions. To accurately describe the effects of disordered tails, it is necessary to incorporate a range of unbiased spatial conformations of the disordered tails into the theory. A comprehensive treatment of disordered histone tails, as well as extending the theory to include multiple histone octamers, is left for future studies. Further details of the generation of the nucleosome conformation(s) are presented in the Supporting Materials. Note that although the distribution of the number density is fixed, the position-dependent volume fraction of the nucleosome is not, because ion condensation can alter the nucleosomal volume.

The term, *E*_VdW_, describes the non-electrostatic attractive van der Waals interactions, representing the hydrophobic interactions of the nucleosome. Since we assumed the nucleosome to be in a frozen state this contribution equates to a constant and does not explicitly need to be considered in the free energy.

The chemical free energy associated with the (de)protonation of the acidic and basic AAs, phosphates, and ion condensation, is described by the term *F*_chem_. The free energy contribution related to the acid-base equilibrium and monovalent ion condensation to the amino acids takes on the following form:^52–54^

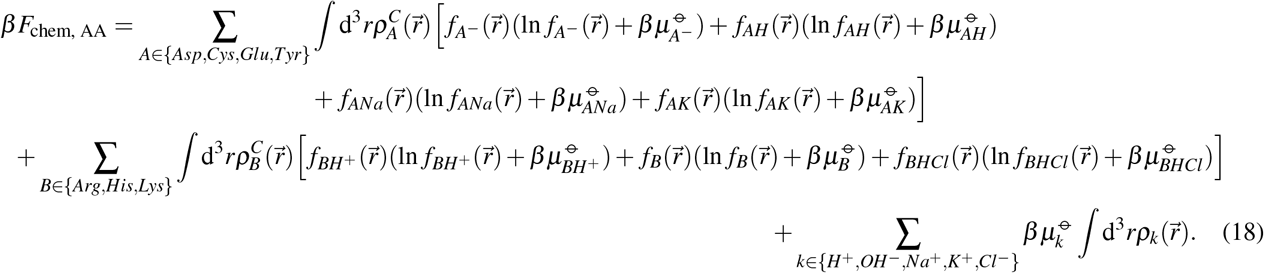

Here, 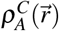 denotes the number density of the chargeable acid centers of acidic amino acids, with A corresponding to Asp, Cys, Glu, or the Tyr amino acid. Similar, 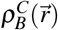 corresponds to the number density of the chargeable centers of the basic amino acids. Namely Arg, His, and 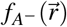 is the fraction of acidic AA residues that are charged or deprotonated at position 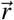, while 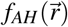 is the fraction of neutral, protonated acids, and 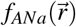 and 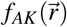 are the fraction of acids that are condensed with Na^+^ or K^+^, respectively. Similarly, 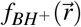 is the fraction of basic amino-acids residues that are charged or protonated at position 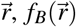 is the fraction of neutral, deprotonated basic amino acid bases, and 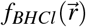 is the fraction of basic amino acids that are condensed with Cl^−^.

In the free energy (Eq. 16), the terms involving 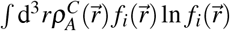 describes the conformational entropy of chemical state *i* of amino acid type *A*. While the terms involving 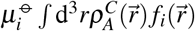 correspond to the standard chemical free energy associated with chemical state *i* of amino acid type *A*. Here 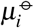 is the standard chemical potential of one amino acid *A* in state *i*. Note it is these chemical potentials that enter into standard reaction Gibbs free energy of the acid-base and condensation reactions (Eqs. 1 through 10).

The chemical free energy contribution associated with the (de)protonation of the DNA-phosphates acids and the ion condensation from divalent ions can be described similarly to that of monovalent ions. However, since we also want to account for the possibility of magnesium bridges, where two phosphates bind simultaneously to a single magnesium ion, 2 P^−^ + Mg^+^ P_2_Mg, a different approach is required. This new approach centers on using the density of phosphate pairs and evaluating whether each pair is bound by a single Mg^2+^ ion or not. By focusing on the density of pairs of phosphate molecules rather than just the density of individual phosphates, we explicitly account for the fact that each *P*_2_*Mg* pair comprises two distinct phosphates. The resulting free energy describing ion-bridging of one Mg^2+^ ion to a pair of phosphates is given by

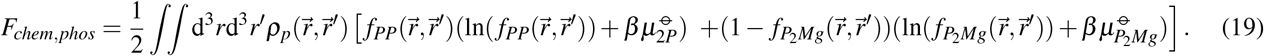

Here *ρ*_*p*_(*r, r*^*′*^)d^3^*r*d^3^*r*^*′*^ is the number of pairs with one phosphate of the pair at volume [*r, r* + *dr*] and the other phosphate located within volume [*r*^*′*^, *r*^*′*^ + *dr*^*′*^]. Thus, only phosphates that are sufficiently close to each other, are to be considered as pairs that can bind to a Mg^2+^ ion. More specifically, the distance between 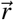 and 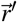 must be less than a cut-off of approximately 0.8*nm*. This distance is comparable to the separation between two phosphates located on the same DNA strand that belongs to adjacent base pairs. The term *ρ*_*p*_(*r, r*^*′*^)d^3^*r*d^3^*r*^*′*^ describes the thermodynamic average pair density function of a phosphate pair located at 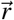 and 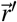.

In free energy of Eq. 19, 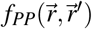 is the fraction of phosphates pairs located at 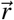 and 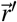 that are both charged and 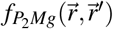 is the fraction of the pairs of phosphates that are bound with one Mg^2+^ ion. The first and third terms describe the conformational entropy of forming an ion bridge, and the third and fourth terms involving *µ*_*P*_ and 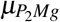 correspond to the standard free energy associated with the charged state and the state where one magnesium is bound to a phosphate pair. The phosphate pair is assumed to be in a completely charged state (*PP)* or in a Mg-bridged state (*P*_2_*M*_*g*_). A generalization of the free energy to include other chemical states of phosphate pairs can be found in the supporting materials. These various chemical states of the pairs include cases such as one phosphate being deprotonated and the other phosphate is protonated, *P*(*PH*) or *(PH*)*P*, or situations where one or both are bound to ions such as potassium, *P*(*PK*), (*PK*)*P* or (*PK*)(*PK*), among others. In total, 25 distinct chemical states for the phosphate pairs are considered, not including the Mg-bridge state (*P*_2_*M*_*g*_). We introduce a fraction denoted as 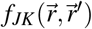 for each chemical state associated with the phosphate pair. The complete free energy, along with further background on its derivation, is presented in the supplementary material, where a more extensive generalization of the free energy discussed here is also provided. Once the fractions of the different chemical states of phosphate pairs, along with those of acidic and basic AAs and their densities, have been determined, the total charge distribution can be calculated.

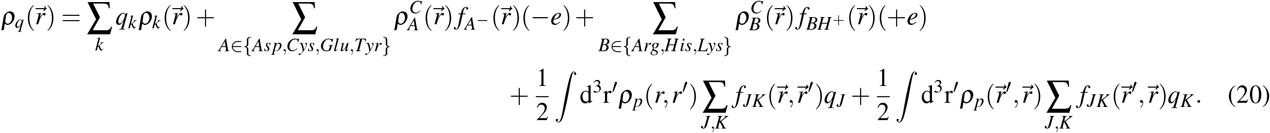

The first term in the equation describes the contribution to the charge density from free mobile ions, with *q*_*k*_ and 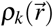 corresponding to the elementary charge and number density of mobile species *k*. Here *k* runs over *k* ∈ {*k H*^+^, *OH*^*−*^, *Na*^+^, *K*^+^, *Mg*^2+^,*Cl*^*−*^. The second and third terms describe the charges of the acidic and basic AAs, while the last two terms relate to the charge of the phosphates. Here *J,K* runs all the chemical states of the phosphate pairs not including the magnesium bridge, i.e., *J, K* ∈ *{P*^*−*^, *PH, PNa, PK, PMg*^+^*}* with 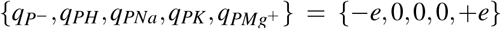 being the charge of that chemical state. A similar term describes the volume fraction contribution of the phosphates.

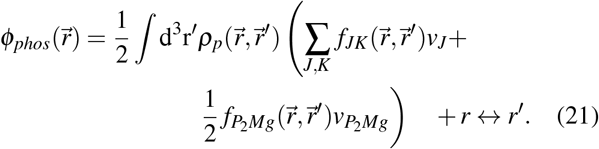

The charge distribution is explicitly represented in *E*_*elect*_, which describes the electrostatic energy contribution to the free energy and couples the charge distribution with the electrostatic potential: 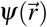 The variation of the total free energy with respect to the electrostatic potential, 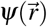, yields a generalized Poisson Equation. It is important to note that, in this Poisson Equation, the charge density and electrostatic potential are replaced by their thermodynamic averages, a consequence of the mean-field approximation as discussed in^53^ and supporting the material. Consequently, fluctuations and short-range electrostatic correlations are not explicitly considered. However, these short-range interactions are implicitly represented through a chemical equilibrium approach, which offers an intuitive way of introducing electrostatic ‘correlations’ and short-range electrostatic interactions at a mean-field level.

To determine the equilibrium structure and charged state of the NCP as well as the density distribution and chemical states of all species the free energy needs to be minimized with respect to the density profiles of all molecular species, the free energy must be minimized with respect to the density profiles of all molecular species, 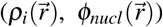, the different chemical states of the amino acids 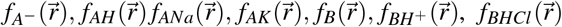 and the chemical states of the phosphates, 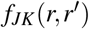 and 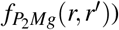, and also with respect of the electrostatic potential 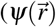.

Minimization of the free energy with respect to the different charged states of the acidic amino acid results in the following set of equations that determine the chemical equilibria:

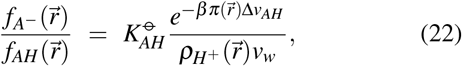

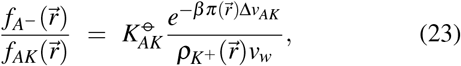

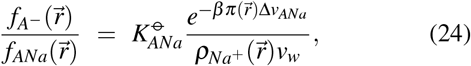

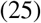

The variable 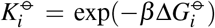 the chemical equilibrium constant, where 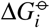 is the standard free energy change for either the acid-base equilibrium reaction of an AA acid or the (dissociation) equilibrium reaction of acid-ion pairs like ANa or AK. The term Δ*v*_*i*_ denotes the volume difference between the products and reactants. Specifically, for the standard free energy of the acid-base equilibrium, AH ⇌ A^−^≡H is given by 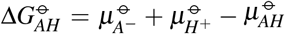 and the volume change is equal to 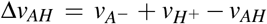. The chemical equilibrium constant 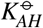 is related to the experimental acid-base equilibrium constant 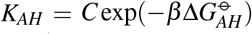 for a monomer in infinitely dilute solution. Here, *C* is a constant required for the consistency of units. The constant *C* arises because the equilibrium constant is expressed in units of the reference solution, which conventionally is chosen to be the 1-molar solution. Therefore, *C* = 1/*N*_*A*_*v*_*w*_, where *N*_*A*_ is Avogadro’s number. Similarly, the chemical equilibrium constant, the standard free energy change for ion condensation reactions, and changes in volume by the reactions for ANa ⇌ A^−^ + Na^+^, and AK ⇌ A^−^ + K^+^ are defined in the same manner as the acid-base equilibrium. It is important to recognize that for both ion condensation and ion bridging, Δ*G*, represents the internal free energy difference between the unbound, solvated phosphate pair and the solvated magnesium state, and the bound, solvated state of the phosphate pair bound magnesium. Therefore, Δ*G* captures the internal free energy change that occurs during ion bridging, implicitly accounting for modifications in the solvation layer as well as shifts in internal electrostatic interactions, without explicitly modeling the hydration forces among water, ions, and charged groups.

Minimizing the free energy with respect to various fractions of (charged) states of the basic group of the different AAs yields similar equations, each involving a specific equilibrium chemical constant related to the protonation or condensation of each basic group. Similarly, minimizing the free energy with respect to the different fractions of phosphate-charged states results in a more complex set of equations, detailed in the supporting information. Note that only in the limit of infinite dilution does the degree of (de)protonation and condensation align with ideal chemical reaction equations. In practice, the fractions of charged states for AAs and phosphates deviate from the ideal solution values because these molecules are not in an ideal solution state. They are in close proximity to each other and interact through electrostatic and excluded/osmotic forces as indicated by locally varying 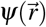 and 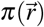, which substantially deviate from ideal conditions. This non-ideal behavior also affects the density of protons and ions, whose local concentrations can differ greatly from the reservoir concentrations. Therefore, chemical interactions are coupled with electrostatic interactions and steric repulsions, resulting in non-ideal solution behavior.

The values of the acid-base chemical equilibrium constants, along with the abundance of AAs and DNA phosphate groups within the nucleosome, are provided in Table S1 of the Supporting information. Experimental data is available for the acid-base equilibrium constants of (weak) AAs. The acid-base constant of DNA phosphate value, a strong acid, is less precisely known but is typically reported as *pK*_*a*_ = 1. The various chemical equilibrium constants associated with ion condensation reactions are far less well established, particularly for Mg-bridging interactions, which are difficult to determine, and to our knowledge, have not been reported. Consequently, we estimate the ion binding energies using a combination of (indirect) experimental observations, theoretical calculations, and past simulations of similar systems.

The free energy minimization that determines the charge state of the NCP results in a system of nonlinear coupled integro-differential equations with unknowns being (1) the osmotic pressure, or Lagrange multipliers that enforce the incompressibility constraints (*E*rep), and (2) the electrostatic potential. Through discretization, these integro-differential equations are converted into a set of non-linear algebraic equations, which can be solved iteratively using standard numerical methods^55^. These equations self-consistently determine the density profiles of all molecular species 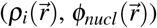, the charged states of the AAs and DNA-phosphates, 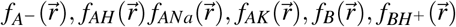, and 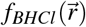 and chemical state of the phosphates, 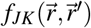 and 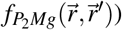 in terms of the lateral pressure 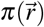 and electrostatic potential 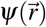. The density of the DNA phosphates, and amino acids as well their charged chemical state as expressed by 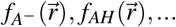 link and couple the chemical interactions with the electrostatic and excluded volume interactions and are coupled with packing constraints. The interplay between chemical, structural, and physical interactions is most clearly represented in the equations for the acid-base and ion-condensation reactions. Therefore, the charge state of all chargeable AAs and phosphates of the nucleosome results from the combined effect of the chemical, electrostatic, and packing or excluded volume interactions. Detailed expressions for the ion, solvent, phosphate, and amino acid densities, along with their chemical states, are provided in the supporting material. Additional information on the numerical methods, including discretization and nucleosome conformation generation, is also available.

## III. RESULTS

### A. Global Effects of Divalent Cations

Here we examine how divalent ion concentrations affect the electrostatic behavior of an NCP with and without histone tails. To characterize the nucleosome’s charging behavior and quantify the effects of varying salt concentrations, we computed the average total of charge of the NCP, denoted as *Q*_*nuc*_. This value is obtained by integrating the local, position-dependent charge density of the AAs and phosphate charge density and summing all contributions. This corresponds to the integration of the last three contributions of the total charge density of Eq. 20

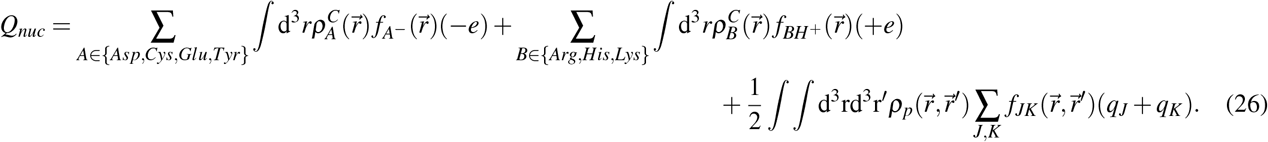

To establish a basis for analyzing the impact of divalent cations on an NCP, we first examined the total charge of nucleosomes with and without histone tails in the presence of increasing concentrations of monovalent potassium ions (K^+^) under physiological conditions, with a pH of 7.4 and 10 mM NaCl, but in the absence of divalent ions (Figure 2). Our findings show that as the concentration of K^+^ increases, the overall nucleosome charge becomes less negative. This effect arises from K^+^ ions binding to the negatively charged phosphate groups, partially neutralizing the nucleosome’s charge. However, the effect of K^+^ ions on nucleosome charge is relatively modest due to the single charge contribution a single K^+^ has. Having characterized the effect of monovalent cations in physiological salt and pH conditions, we now turn our attention to examining how divalent cations influence nucleosome charge and stability.

**FIG. 1.**
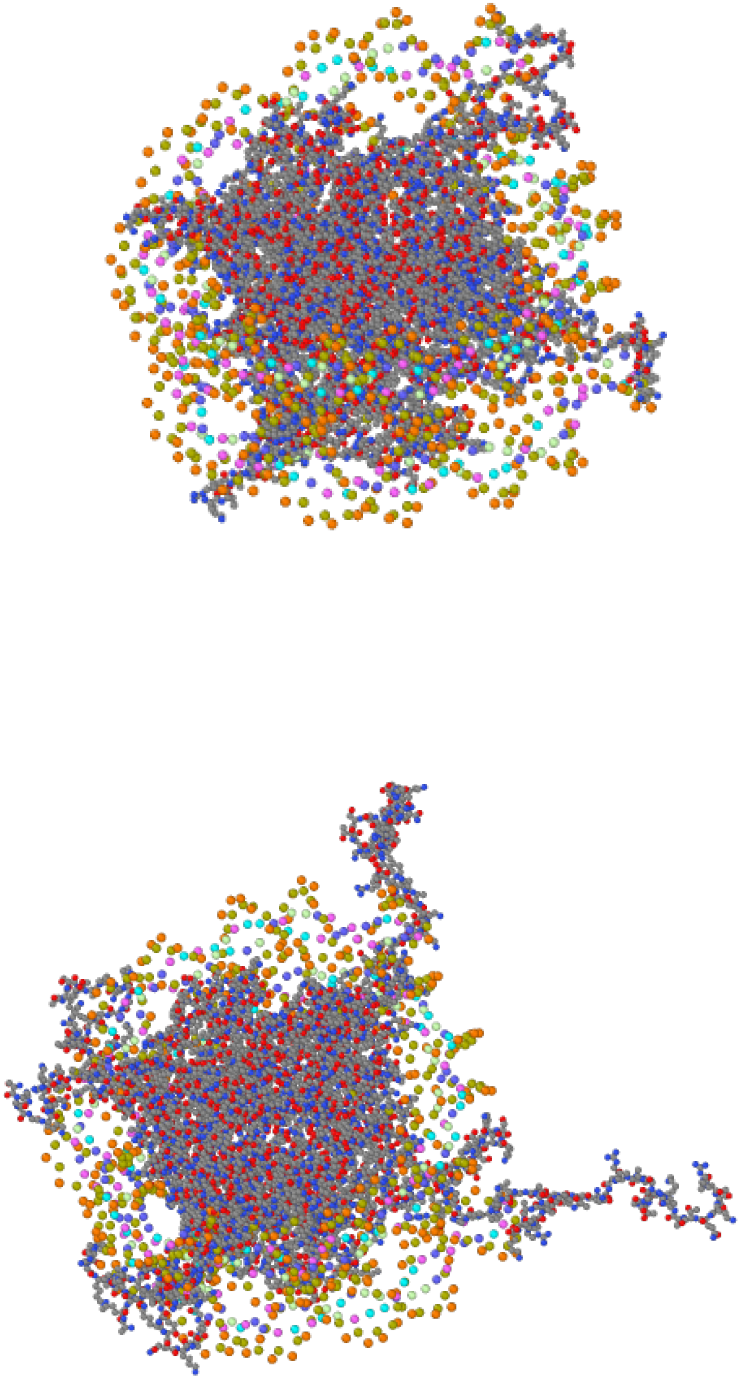
Molecular representation of a single nucleosome without (top) and with disordered histone tails (bottom). Observe that our system explicitly contains the most prevalent intranuclear ions, including Na^+^, K^+^, Mg^2+^, and Cl^−^ as well as water, OH^−^ and H^+^, to account for charge regulation and acid-base equilibrium as well as ion-condensation effects.

**FIG. 2.**
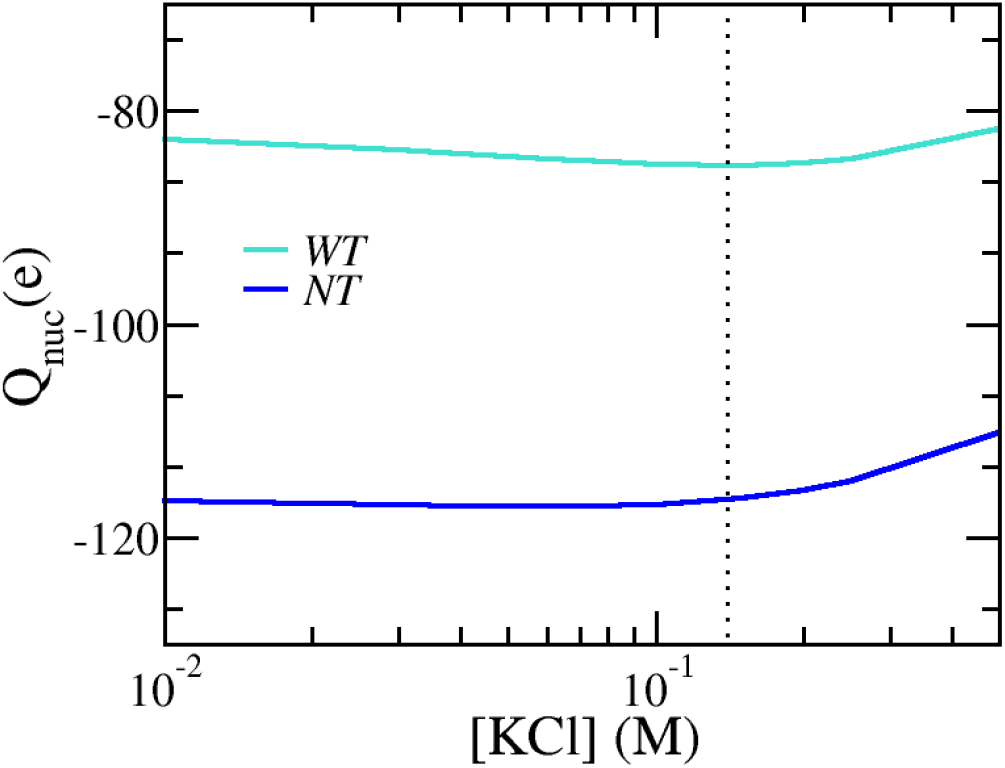
Total charge of a nucleosome with (WT) and without (NT) tails as a function of monovalent K^+^ concentration, in physiological conditions without divalent cations (pH = 7.4, [NaCl] = 10 mM). The vertical dotted line at 140 mM marks the physiological concentration of K^+^.

Figure 3 shows the total nucleosome charge as a function of divalent MgCl_2_ concentration. The pH is set to 7.4, corresponding to that of a physiological solution. The concentra-tion of monovalent KCl and NaCl salts are set to the physiological values of 140 mM and 10 mM respectively. We varied MgCl_2_ across a wide range of Mg^2+^ concentrations, including those relevant to physiological conditions, as indicated by the vertical dotted lines and the inset. Very low Mg^2+^ levels were also considered to explain the observed charging behavior of the nucleosome.

**FIG. 3.**
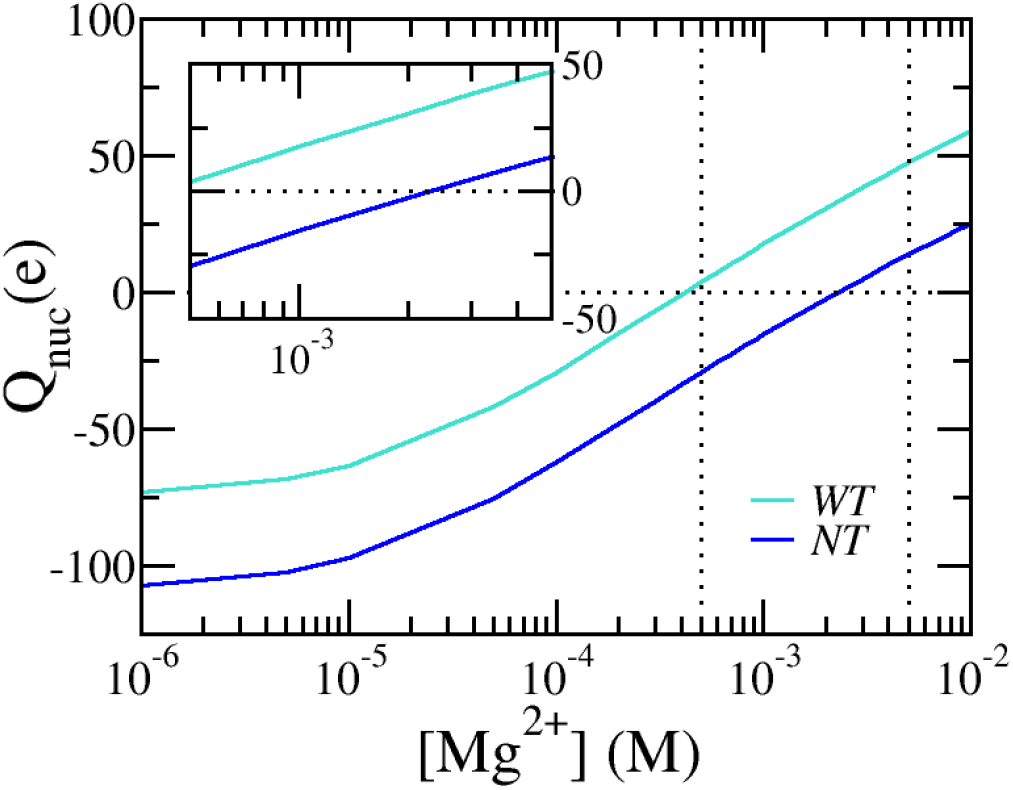
The total nucleosome charge with (WT) and without tails (NT) as a function of divalent Mg^2+^ concentration for physiological conditions: pH=7.4, [KCl]=140 mM, and [NaCl]=10 mM. The vertical dotted lines at 0.5 mM and 50 mM mark the physiological range for magnesium, while the horizontal dotted line at 0 marks the charge inversion point.

Determining the concentrations of nuclear ions in cells, both monovalent and divalent, is very challenging. For example, patch-clamp techniques, commonly used in electrophysiology to measure ionic currents, infer ionic concentrations from the membrane potential, but cannot distinguish between cytosolic and nuclear concentrations or different ion types. Fluorescence microscopy can distinguish different ion types but cannot measure absolute concentrations^56,57^ Techniques such as electron probe X-ray microanalysis, ^31^P-NMR, selective Mg^2+^-electrode, and fluorescent indicators, have de-termined total Mg^2+^ concentrations to be approximately 20 mM inside many (mammalian) cells^58,59^. Free magnesium concentrations of 0.8 − 1.2 mM have been reported for the cytoplasm and extracellular matrix^58^, but free magnesium concentrations inside the nucleus are not well documented. The observed difference between cytoplasmic and nuclear magnesium levels suggests that most magnesium ions in the nucleus are bound to proteins, chromatin, or phospholipids in the nuclear envelope, assuming there is no chemical gradient of free magnesium between the nucleus and cytosol. Based on these observations, we set the physiological range for magnesium from 0.5*mM* to 50 *mM*.

It is important to emphasize that the pH and salt concentration in Figures 3 and 2, and throughout the paper refer to the proton and ionic concentration of the reservoir bath with which the nucleosome system is in equilibrium. The reservoir concentration establishes the chemical potentials for water, protons, hydroxyl ions, and other ionic species which are constant throughout the system according to thermodynamic equilibrium. However, the local position-dependent ion and proton concentrations can differ significantly from their concentrations in the reservoir, or free concentrations, as will be explained later.

Fig.3 demonstrates a strong dependence on divalent Mg^2+^ salt concentrations. At low Mg^2+^ concentrations, the nucleosome is highly negatively charged. However, as Mg^2+^ concentrations increase starting from the submillimolar ranger, there is a much more profound effect than that caused by monovalent ions in the net negative charge. For example, at [Mg^2+^]=0.1 mM the net nucleosome charge is *Q*_*nuc*_ = − 62.3*e* which decreases to *Q*_*nuc*_ = − 29.3*e* at [Mg^2+^]=0.5 mM and further drops to *Q*_*nuc*_ = − 15.6*e* for [Mg^2+^]=1 mM. This represents a three-fold reduction in nucleosome charge as the Mg^2+^ concentration increases from 0.5 mM to 1 mM. At concentrations around [Mg^2+^]=2.5 mM, the effective nucleosome charge neutralizes, and at higher Mg^2+^ concentrations, the effective nucleosome charge becomes net positively charged, indicating a reversal in charge. Even within physiological ranges, a significant shift in the nucleosome’s charge state is observed as magnesium concentrations increase, rising from approximately − 29*e* to + 13*e*. The same trend is similarly observed in an NCP with disordered histone tails.

To gain further insight into the charging behavior of the nucleosome and its strong dependence on Mg^2+^, we present Figs. 4, 5 and 6. These results pertain to a nucleosome without tails. Figure 4 separately shows the negative charge of the phosphates and the net positive charge of the histone octamer separately, along with their sum, representing the total nucleosome charge: *Q*_*nuc*_ = *Q*_*hist*_ + *Q*_*phos*_. Given the nucleosome composition and assuming ideal acid-base equilibrium, we would expect that all 292 phosphates of the nucleosome would deprotonate, leading to a total phosphate charge of *Q*_*phos*_ = − 292*e*. The core histone octamer contains 92 lysines and 82 arginines, the most abundant basic AAs contributing to the positive charge. When including the histone tails, there are an additional 34 lysines and 12 arginines that further contribute to the charge and lead to an upward shift when compared to a nucleosome without tails. Since the number of DNA phosphates far exceeds the number of basic AAs, the nucleosome, with and without histone tails, is imbued with a large net negative charge. Besides lysine and arginine, there is also a considerable amount of glutamic (48) and aspartic acid (22). Considering all AA, the total ideal net charge of the amino acids would be *Q*_*hist*_ = +104*e*. This would result in a large net negative charge of *Q*_*nuc*_ = − 187.6*e* for the nucleosome without tails. A complete list of chargeable AAs and phosphates, along with their *pK*_*a*_ values is provided in the supporting materials.

**FIG. 4.**
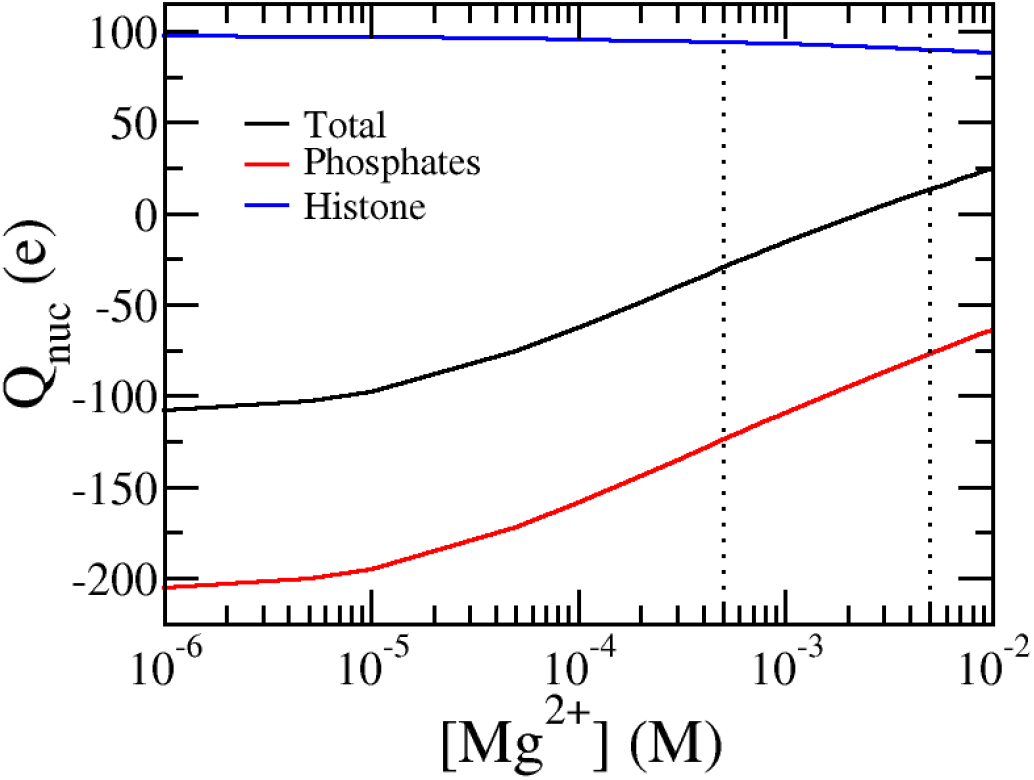
The net positive charge of the phosphates, the charge of the histone octamer, and the total effective charges of the nucleosome presented as a function of Mg-concentration for a nucleosome without disordered tails. As Mg^2+^ concentration increases, the total nucleosome charge becomes less negative, reflecting increased charge screening and partial neutralization of the phosphate backbone. The vertical dotted lines at 0.5 mM and 50 mM mark the physiological range for magnesium. The conditions are pH=7.4, [KCl]=140 mM, and [NaCl]=10 mM.

**FIG. 5.**
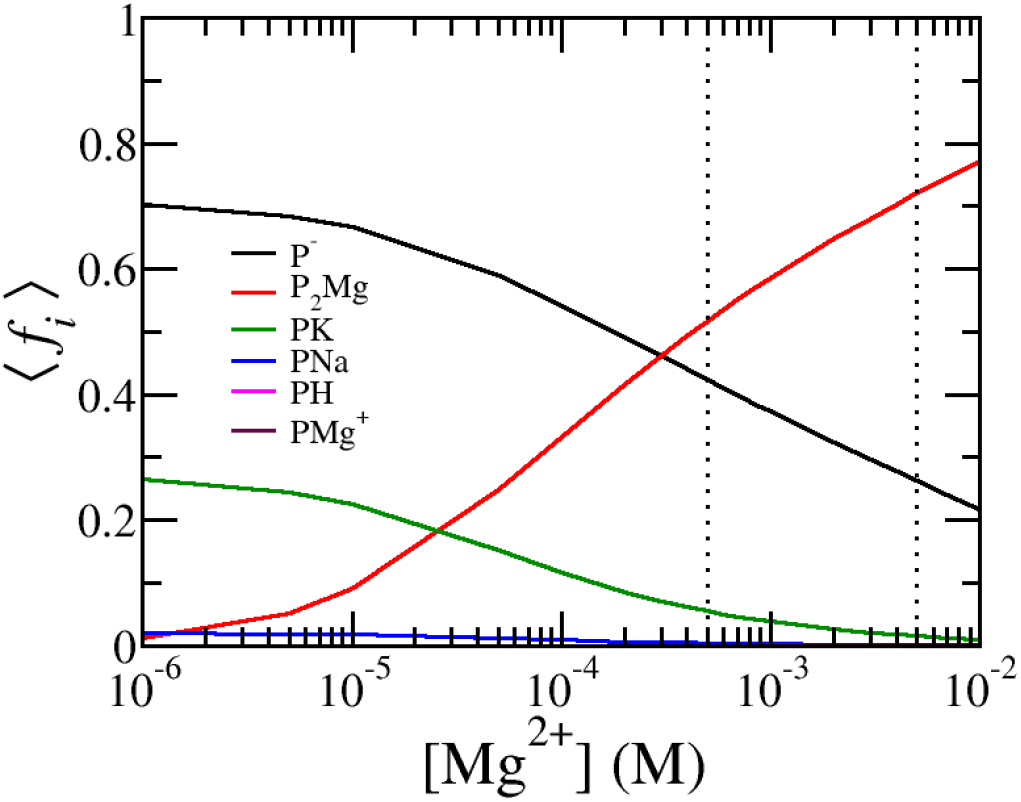
The average fraction of chemical states of the phosphates as a function of Mg^2+^ concentration. The deprotonated (charged) state is labeled *P*^*−*^, the magnesium bridge (*P*_2_Mg), the phosphates bound with K^+^, Na^+^, or Mg^2+^ are denoted as *PK, P*Na, and *P*Mg^+^, and the protonated state is labeled *PH*. As Mg^2+^ concentration increases, magnesium dominates the binding and significantly influences the chemical states of the phosphates, particularly in the physiological range (0.5 mM to 50 mM), as seen by the increased fraction of *P*_2_Mg and *P*Mg^+^. The conditions are pH = 7.4, [KCl] = 140 mM, and [NaCl] = 10 mM.

**FIG. 6.**
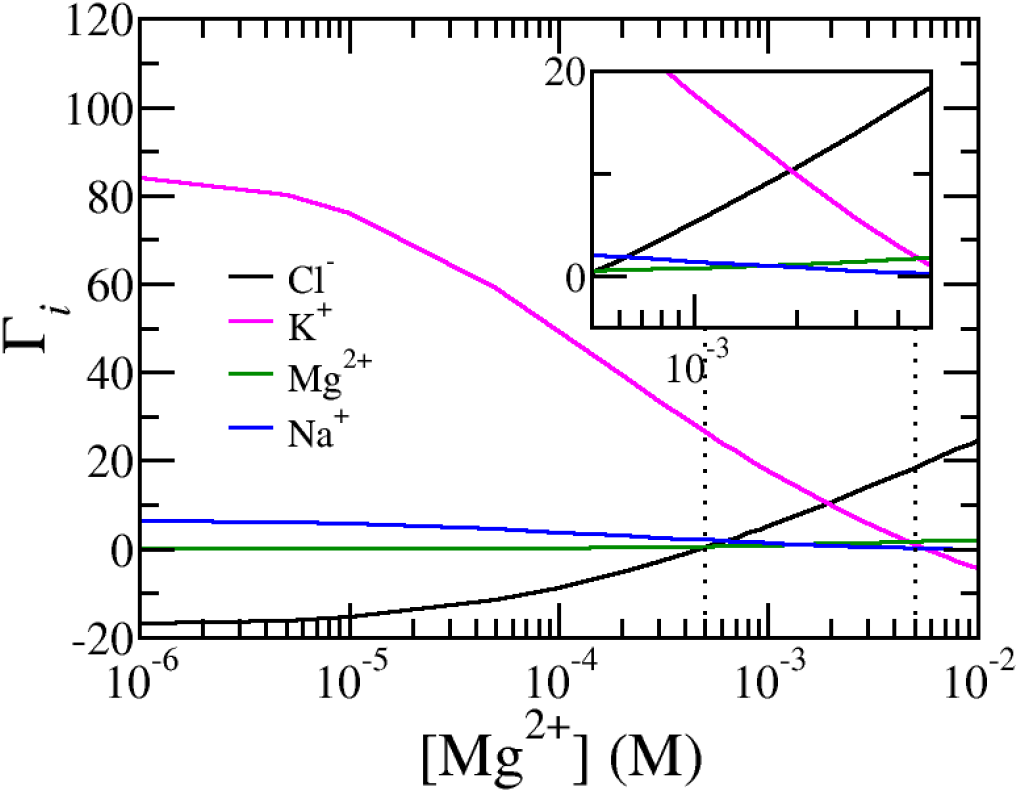
The ion excess of free K^+^,Na^+^,Mg^2+^ and Cl^−^ ions as function of Mg^2+^ concentration for pH=7.4, [KCl]=140 mM, and [NaCl]=10 mM. As the nuclesome’s charge flips from positive to negative chloride ions replace potassium ions as counterions. The vertical dotted lines at 0.5 mM and 50 mM mark the physiological range for magnesium.

### B. Mechanisms of charge regulation

Fig.4 shows that both the number of phosphate charges and the net nucleosome charge are significantly lower than the expected ideal value, c.f. the total ideal charge of -187e to actual net charge of -15.6e at 1mM of magnesium. We also observe that the net negative charge of the phosphates decreases substantially with increasing Mg^2+^ concentration, while the net positive charge of the histone octamer shows only a slight reduction. Since there are far more phosphates than positively charged AAs, the nucleosome remains net negatively charged for magnesium concentrations up to 2.5 mM. Beyond this concentration, the net nucleosome charge reverses in sign.

Due to the asymmetry between negative and positive charges, the charges cannot be perfectly balanced, and if ideally charged, the nucleosome would experience large electrostatic repulsions between like charges. To mitigate this excess repulsion, the nucleosome system can employ several mechanisms: counterion confinement, ion condensation, and ion pairing, or shifting of the acid-base equilibrium of AAs and phosphates. Counterion confinement involves attracting counterions from the reservoir to enhance electrostatic screening. This process reduces enthalpic electrostatic repulsions by localizing counterions near oppositely charged regions, though it comes with an entropic cost due to the loss of translational entropy of the counterions. Electrostatic repulsion can also be reduced by decreasing the net charge of the chargeable AAs and phosphates. This can be accomplished through ion condensation, where oppositely charged ions bind to chargeable moieties, or by shifting the acid-base equilibrium toward the neutral state of the AAs or phosphates. The latter occurs at the cost of standard chemical free energy of the acid-base reaction. 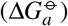. Ion pairing, on the other hand, is accompanied by a free energy gain which is the negative of the dissociation free energy 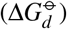.

The primary mechanism by which the number of phosphate charges is reduced is through ion pairing with K^+^ and the formation of magnesium bridges as illustrated in Fig.5, which shows the chemical state of the phosphates as a function of Mg^2+^ concentration. Here *f*_*i*_ represents the average fraction of phosphates in chemical state *i* normalized by the total number of phosphates. The various chemical states include (de)protonation, condensation with K^+^, Na^+^ or Mg^2+^, and the magnesium bridge state (P_2_Mg). These average fractions can be obtained by integrating the local fraction of the phosphate pairs in chemical state *i* weighted with the local phosphate pair density. For example, the fraction of phosphate bond with K^+^ is given by

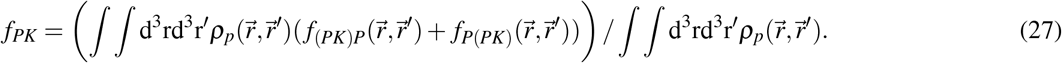

The other fractions are defined similarly and their definitions are listed in the supporting materials. Here, Fig. 5 shows that at low magnesium concentrations, about 70% of the phosphates are charged while 30% of the phosphates are condensed with potassium. A small fraction of phosphates is also bound with sodium, but because, the concentration of potassium is much higher, the likelihood of sodium condensation is much lower.

As Mg^2+^ concentrations increase, we observed magnesium-ion bridges begin to form, replacing the potassium-phosphate ion pairs. At physiological and high Mg^2+^ concentration, the P_2_Mg bridge becomes the dominant chemical state of the phosphates. At [Mg^2+^]=1 mM the fraction of phosphates that are part of an P_2_Mg bridge equals 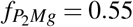 and reaches a value of 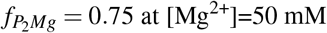 at [Mg^2+^]=50 mM. Divalent Mg bridging is more favorable than monovalent ion pairing because one (P_2_Mg) bridge electrostatically neutralizes two phosphate charges, whereas one monovalent ion neutralizes only one phosphate. To achieve a similar reduction in electrostatic interactions, two monovalent potassium ions would need to pair, but this would in roughly twice the loss of translation entropy. Therefore, forming Mg-ion bridges is energetically more favorable than monovalent ion pairing. As Mg^2+^ concentration increases, the nucleosome exhibits a strong tendency to form *P*_2_*Mg* magenisum bridges, leading to a reduction in its effective charge.

So far, the charging behavior of the nucleosome has been explained primarily in terms of phosphate charging, which plays a key role in regulating the nucleosome’s charge. However, this does not suggest that ion bridging of Mg^2+^ and phosphates is the sole mechanism at work. Other mechanisms, such as counterion confinement and charge regulation through shifts in the acid-base equilibrium mentioned above, also contribute to mitigating electrostatic interactions. To demonstrate the importance of these additional mechanisms, Fig. 6 presents the ion excess of the ions surrounding the nucleosome. Ion excess for a given ion type is defined as the difference between the local, free ion number density and the ion number density of the reservoir integrated over the entire space.

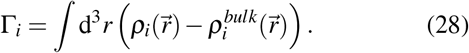

Due to charge neutrality, the sum of the ion excesses for all charge mobile species, multiplied by its charge, equals the negative of the total nucleosome charge: ∑_*k*_ Γ_*k*_*q*_*k*_ = − Q_nuc_. Thus, the ion excess measures the composition of the counter ion cloud surrounding the nucleosome.

For low magnesium concentrations, the nucleosome is surrounded by a counter ion cloud, which contains an excess of 80 potassium ions, while chloride, a co-ion, is severely depleted from the region surrounding the nucleosome. This substantial excess of potassium ions indicates a high concentration of positively charged ions near the nucleosome, forming a double layer that screens electrostatic interactions. The large value of the ion excess indicates that the electrostatic interactions are strong. The magnitude of this potassium ion excess roughly corresponds to the nucleosome’s net charge. As magnesium concentrations increase, more (P_2_Mg) bridges are formed, reducing the number of negatively charged phosphates. Consequently, fewer counterions are needed to screen the electrostatic interactions, leading to a decrease in potassium ion excess. A visual representation of the reduction in the electrostatic potential can be found in the supporting materials, Fig. S1. Sodium ions, which also contribute positively to the counterion cloud, show a similar behavior to potassium ions but are present in much lower concentrations (10 mM of Na^+^ versus 140 mM of K^+^).At high magnesium concentrations, exceeding physiological levels, the excess of potassium ions becomes slightly negative. This is because the nucleosome’s net charge switches from negative to positive at these high magnesium levels. In this scenario, negatively charged chloride ions replace potassium ions as counterions, balancing the positively charged nucleosome. Chloride ions, which are less abundant than potassium ions, exhibit the opposite trend: they are initially depleted at low magnesium concentrations but become more confined and in excess as magnesium levels rise. Notably, chloride ions are already present in excess even when the nucleosome’s net charge remains negative. This behavior occurs within physiological ranges, as shown in the inset in Fig. 6 where large variations in ion excess as a function of magnesium levels are observed among all four ions in the system, especially between potassium and chloride.

To explain this seemingly surprising result, it is important to realize that the ion excess, total charge, and fraction of chemical phosphates states, defined by Eqs. 26-28 and plotted in Figs. 4,5, and 6 relate to spatial averages. If the distribution of amino acids (AAs) and phosphates were uniform, these global averages would accurately reflect the local charging behavior. However, the distribution of AAs and phosphates is not uniform. As seen in Fig. 1, DNA phosphates are wrapped around the ‘equator’ of the histone core, while different AAs are distributed unevenly within the histone octamers. The various histone proteins have distinct AA compositions, with positively and negatively charged amino acids clustered rather than evenly spread. This results in localized regions with higher concentrations of acidic or basic AAs, leading to areas of net negative or positive charge. Notably, the negatively charged glutamic and aspartic acids form regions of negative charge, often referred to as acidic patches.

To explore how the distribution of AAs and phosphate affect nucleosome charging, we present Fig. 7 which illustrates the distribution of the electrostatic potential, the local proton concentration, the local potassium concentration as well as the total volume fraction of the nucleosome with and without tails. The total volume fraction of the nucleosome exhibits significant variation: histone proteins appear as dense regions with volume fractions up to 90%, whereas the DNA backbone surrounding the core is much less dense, with a volume fraction of approximately 20%. The chosen spatial resolution of *δ*=0.65 nm, enables us to clearly identify major features of the nucleosome, such as the wrapped DNA and the histone-disordered tails. Minor features of the nucleosome are also visible, including the density drop in the center of the histone core as well as the helical twist of the wrapped DNA. The latter is most clearly visible in the movie of the electrostatic potential, which allows the electrostatic potential to be viewed from different viewpoints. The movie is found in the supporting materials.

**FIG. 7.**
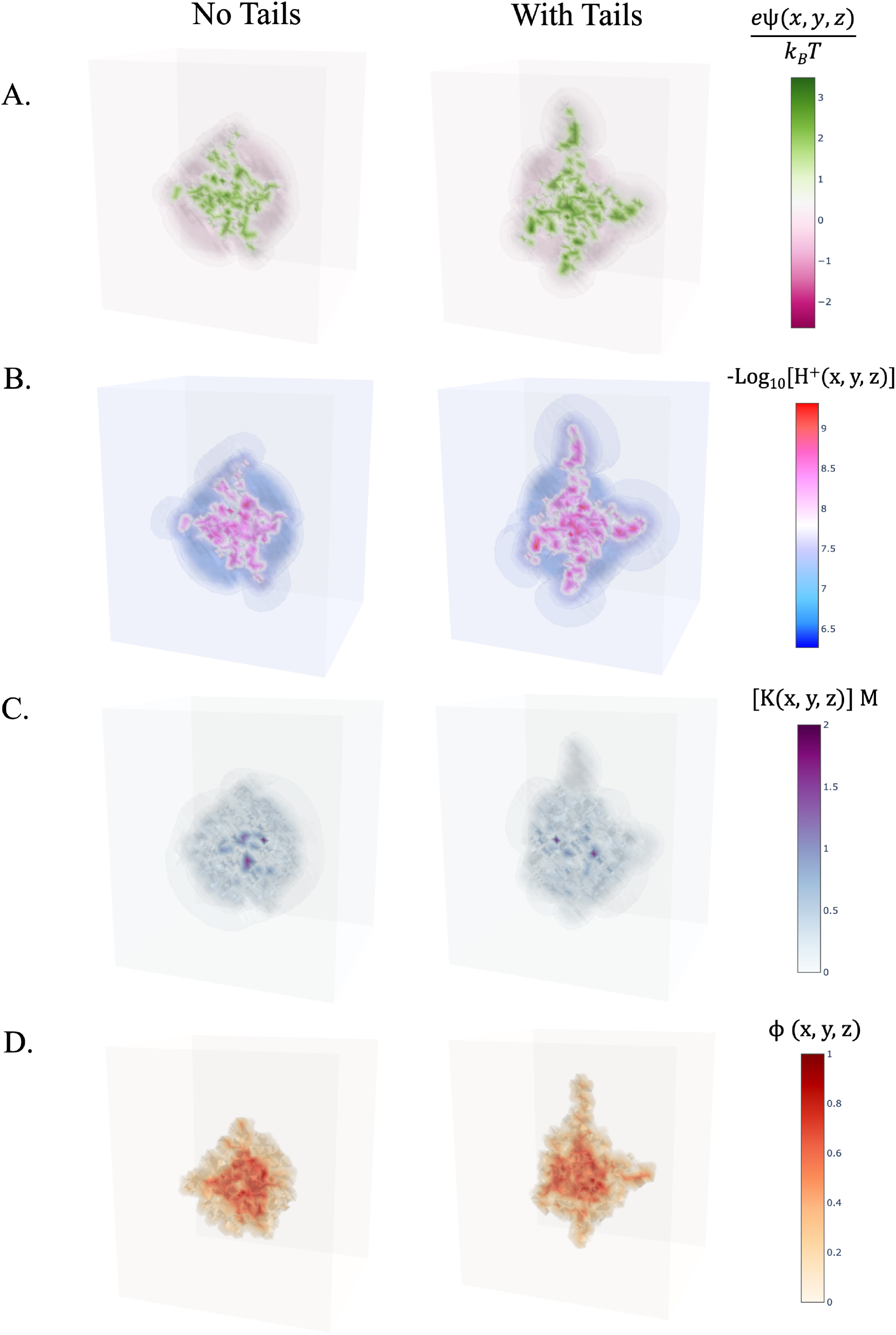
Distribution of (a) electrostatic potential, (b) local position-dependent proton concentration (c) local concentration of KCl, (d) local total volume fraction of the nucleosome at physiological conditions. Namely pH=7.4, [KCl]=140 mM, [NaCl]=10 mM, and [MgCl_2_]= 1 mM. Each plot compares nucleosomes without tails (left) and with tails (right), highlighting the increased heterogeneity within the structure. The graphs, generated using the Plotly-Python package^60^, are semi-transparent to improve visualization and contrast the distribution against the reservoir values. (Opacity = 0.1). The height, width, and depth are respectively 19.5 nm, 22.75 nm, and 26 nm for all graphs.

In the electrostatic potential 3D plot, magenta regions indicate negative electrostatic potential values which correspond to the location of phosphates or acidic AAs. Green regions, on the other hand, represent positive electrostatic potential values highlighting the presence of basic AAs. Details on the corresponding charge densities are available in the supporting materials. The negative electrostatic potential around the core of the nucleosome is indicative of phosphate charges. Similarly, the major H3 histone tail, rich in lysines, is visible as a region of positive electrostatic potential. The electrostatic potential within the core is heterogeneous, with areas of high positive potential alternating with regions of negative potential.

The heterogeneity of the electrostatic potential mirrors the variation in local K^+^ concentration. Here, the ‘darker’ regions correspond to local increase of K^+^ concentration, while the ‘lighter’ color regions correspond to regions from which K^+^ ions are depleted. The regions with negative electrostatic potential, where DNA-phosphates are present, coincide with areas of high K^+^ concentration where K^+^ ions are concentrated to counteract the local negative electrostatic potential. This localized ion confinement can lead to large gradients in the K^+^ concentrations with maximum levels exceeding 2 M, roughly 15 higher than the reservoir concentration of 140 mM. Conversely, ‘lighter’ regions in the K^+^ distribution reflect the lower and even depleted K^+^ concentrations. The lowest K^+^ concentration observed here is 3 mM. Notice also a dark spot inside the histone corresponding to the local region of acidic AAs. The ‘halo’ that surrounds the H3-tail represents a K^+^ ion-depleted zone, correlating with the positive electrostatic potential and charge of the H3 tail.

The distribution of chloride ions exhibits behavior opposite to that of potassium ions. Chloride ions are depleted from areas of local negative electrostatic potential and nucleosome charge, while they accumulate in regions with net positive electrostatic potential and positive charge. This behavior accounts for the seemingly unexpected observation that the ion excess of chloride ions is positive, even when the nucleosome’s net charge remains negative. At low magnesium concentrations, chloride ions are depleted from the negatively charged phosphates, leading to a local negative ion excess. Since phosphates are more numerous than positively charged amino acids, this results in a global negative ion excess. However, as magnesium-phosphate bridges form and neutralize phosphate charges, the depletion of chloride ions from these regions decreases. Nonetheless, chloride ions remain concentrated around positively charged lysines and arginines. This explains the overall pattern of chloride ion excess observed in Fig. 6.

The position-dependent variation is also evident in the local proton concentration. We define a local pH as the negative logarithm of the proton concentration. Significant pH fluctuations are observed, with local pH values inside the core rising to 8–8.5, even higher than in the tail regions. In contrast, the pH in the DNA region drops to approximately 6.5.

## C. Effect of amino acids

Finally, it is important to note that the nucleosome’s charged state is also influenced by acid-base equilibrium and ion pairing of the histone core’s AAs. Fig. 4 shows that the global net charge of all AAs is only slightly affected by changes in magnesium levels. However, this does not imply ideal behavior for individual amino acids. Based on the pKa values, one would expect AAs to be fully charged at physiological pH. For instance, the lysine and arginine AAs would ideally be fully protonated and charged with an ideal average degree of charge 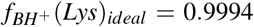 and 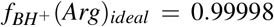 respectively. Considering ideal ion condensation, these values only slightly decrease to 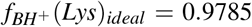 and 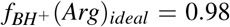. However, our findings show that the actual average degrees of charge are 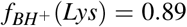 and 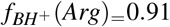. A 11% and 10% reduction in charge respectively. This charged reduction is mostly due to chlorine ion condensation but also to a lesser extent by shifting the acid-base equilibrium. E.g. *f*_*B*_(*Lys*) = 0.1 and *f*_*BHCl*_(*Lys*) = 0.088. Ion condensation results in a loss of translational entropy, while shifting the acid-base equilibrium incurs chemical work, which is considerable for lysine and arginine at physiological pH. To reduce electrostatic repulsion, approximately 9% of lysines and 7% of arginines condense ions. However, this effect is less significant compared to charge reduction through monovalent potassium-phosphate condensation and magnesium-phosphate bridging. This is because monovalent ion condensation is less effective than divalent ion bridging, and many lysines and arginines are ‘buried’ within the histone core, leading to greater excluded volume or osmotic interactions compared to ions in the less dense DNA regions of the nucleosome.

At the same time, the most abundant acidic amino acids, glutamic and aspartic acid, also show reduced charge and deviate from ideal conditions. Ideally, these acids would be fully deprotonated, with 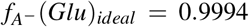 and 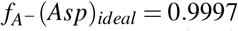. However, their actual average degree of charge are 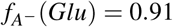 and 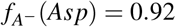 respectively. Since both the most common basic and acidic amino acids experience similar reductions in their charged states, the overall net charge of the histone octamer remains relatively stable despite changes in magnesium concentration. While other AA in the nucleosome core also contribute, because they are less abundant in the system, their contribution is limited. A more detailed discussion on the contribution of these AA can be found in the supplementary material.

The chemical work required to shift the acid-base equilibrium of lysine, arginine, aspartic acid, or glutamic acid, the most abundant amino acids in the histone octamer, is substantial at physiological pH, making the nucleosome’s charge state relatively stable against pH changes. As seen in Fig. 8, significant changes in net nucleosome charge only occur at pH levels well below or above the physiological range. Below pH 6, histone amino acids gain positive charge, while glutamic and aspartic acids lose negative charge, increasing the net positive charge of the histone core and reducing the nucleosome’s overall charge. Above a pH 9, tyrosines deprotonate and lysines lose charge, leading to a reduction in nucleosome charge. Note that pH levels below 6 and above 8 are outside physiologically relevant ranges. Additional results concerning pH-dependence of the charge can be found in Fig S2 of the Supporting Materials.

**FIG. 8.**
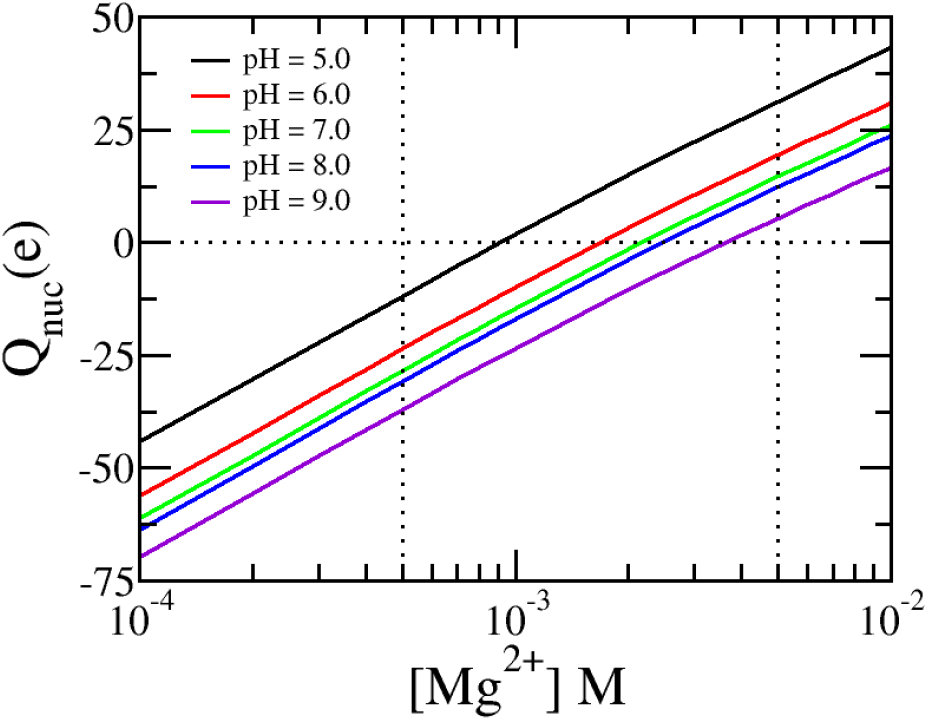
The total nucleosome charge of a nucleosome without tails as a function of divalent Mg^2+^ concentration at different pH values. The vertical dotted lines at 0.5 mM and 50 mM mark the physiological range for magnesium. Remaining conditions: [KCl]=140 mM, and [NaCl]=10 mM.

## IV. DISCUSSION AND CONCLUDING REMARKS

Our main finding is that ionic strength, solution pH, and in particular, divalent magnesium levels exhibit a very large effect on the charged state of the nucleosome, resulting in qualitative and quantitative changes. While nucleosome core particles (NCPs) are naively expected to carry a strong net negative charge based on their composition, we demonstrate that the presence of magnesium-phosphate bridges greatly reduces this net charge. The nucleosome’s charge can switch from net positive to net negative depending on the magnesium concentration in the solution. This charge state arises from a delicate balance involving the chemical dissociation equilibrium and ion condensation of AAs, ion bridging between phosphates and magnesium, electrostatic interactions among nucleosome charges, and the effects mediated by mobile ions and their translational entropy. Consequently, the charging of a single nucleosome is highly non-ideal. Although the net charge of an NCP is significantly reduced, this does not imply weak electrostatic interactions. These interactions remain substantial due to the large local variation of the nucleosome’s charge density. This leads to heterogeneous behavior of the electrostatic potential, ion distribution, and the local proton concentration. Ion and proton distributions also deviate considerably from ideal solution values. Local proton concentration can even vary by more than an order magnitude. These effects are enhanced with the additional lysines that are found when histone tails are added. The predicted heterogeneity highlights the necessity of considering molecular details—such as ion bridging between DNA phosphates and magnesium, and the acid-base equilibrium of amino acids—to fully understand the nature and magnitude of the nucleosome’s charge and its electrostatic interactions under physiologically relevant conditions.

Experiments have shown that NCPs precipitate in solutions containing 2 mM or more of Mg^2+^ and redissolve in concentrations that exceed 50 mM^13,15^. Likewise, experimentally measured electrophoretic mobility of NCPs reverses its sign with increasing multivalent ions, indicative of charge inversion^13^. The present calculations support these experimental findings, given that in millimolar ranges of magnesium the electrostatic potential and net charge drops. The charge neutralizes and reserves sign at elevated Mg^2+^ levels. Observe that our current calculations pertain to a single nucleosome in solution. Thus, we cannot directly compute its phase behavior nor infer the limit of stability against precipitation. However, the reduction of charge and electrostatic potential is a very strong signal in support of the precipitation of charged NCPs. This observation aligns with experiments and simulations, which showed the compaction of nucleosome arrays and chromatin as a function of Mg^2+^ concentration^11,16^. Recent MD simulations also show the clustering of NCPs in the presence of Mg^2+^. Our findings here show that the strong structural Mg-dependence of NCPs and nucleosome arrays is linked with magnesium-phosphate bridging.^30,41^

It needs to be mentioned that all observables such as the charge, isoelectric point, and the strength of the electrostatic potential depend on the choice of various chemical equilibrium constants for acid-base reactions, which are well-documented in the literature^61^, and the ion condensation reactions, which are less well known. The binding energy of potassium and sodium ions to DNA phosphates was set to 3 *k*_*B*_*T*, as this value resulted in an ion condensation degree comparable to the established charge on DNA in simulations. Furthermore, to simplify our estimation, we assumed potassium and sodium ion condensation reactions with all chargeable AAs to have a binding energy of Δ*G*_*bin*_ = − 3*k*_*B*_*T* or equivalently a dissociation constant of *pK*_*d*_ = − 0.444 M. The effect of varying this binding free energy, in the absence of magnesium, on the net charge and ion excess is detailed in Fig. S3 of the supporting material. Our analysis shows that ‘weak’ binding values (1-3 *k*_*B*_*T*) yield similar charging behaviors, while ‘stronger’ binding values (4-5 *k*_*B*_*T*) lead to an increase of the net charge as function monovalent ion concentration which we consider non-physiological. Further details are available in the supporting material. We also set Δ*G*_*dis*_(*PMg*^+^) = − 1.5*k*_*B*_*T* and Δ*G*_*dis*_(*P*_2_*Mg*) = 10*k*_*B*_*T* or equivalently a dissociation constant of *pK*_*d*_(*PMg*^+^) = −1 M and *pK*_*d*_(*P*_2_*Mg*) = 2.5 M respectively. This results in charge neutralization at millimolar concentrations, consistent with experimental observations. Considering, larger ion bridging energies results in a shift of the isoelectric point to unphysiological ranges of magnesium concentration (See Fig. S4 Supporting Materials).

Although quantitative differences arise from variations in the ion binding constants, the results remain robust, with the qualitative charging behavior showing consistent trends regardless of the specific values chosen. A future direction could involve refining these binding constants by calculating the phase diagram of NPC solutions and calibrating it against experimental phase diagrams. Such an approach would allow for a detailed examination of ion-specific effects of monovalent ions on nucleosome charging, which is particularly relevant as experiments suggest K^+^ and Na^+^ to have different effects on nucleosome structure, stability, and their interaction with proteins.^49,62^

Experimental data on the localization of proteins within chromatin, obtained through immunofluorescent staining and confocal scanning laser microscopy, revealed significant Mg^2+^ induced changes in protein distribution that increase with higher magnesium concentration.^63^ The sensitivity of the electrostatic potential to Mg^2+^in millimolar range, along with differences in the NCP changing with and without disordered tails, provides insight into how magnesium levels can influence the degree of nuclear protein association with chromatin. This modulation of nucleosome charge by Mg^2+^ not only affects the spatial organization of nuclear proteins within chromatin but may also impact their enzymatic activity. Since local proton concentrations, or local pH values, which control enzymatic function, exhibit significant spatial variation around the nucleosome, changes in nucleosome charging could play a crucial role in regulating nuclear protein activity.

Additionally, in our investigation, we found that a substantial number of magnesium ions are bound to the phosphates of the NCP at physiological magnesium concentrations. At 1 mM of free magnesium, approximately 50% of the DNA phosphates are bound to magnesium. Assuming similar binding levels occur in the denser nucleosome chains of chromatin, we estimate that bound magnesium concentrations in chromatin could reach 20 mM when free magnesium is 1 mM. This estimate is based on a high nucleosome concentration of 200 *µ*M in chromatin, with around 200 DNA base pairs per nucleosome. This calculation indicates that most of the magnesium ions inside the nucleus are bound rather than free. This suggests that the nucleosome-bound chromatin chromatin may act as a magnesium reservoir or buffer. If free Mg^2+^ levels decrease, nucleosomes would release magnesium ions, and if free Mg^2+^ increases, they would bind more. This buffering capacity could influence chromatin’s structural response to fluctuations in magnesium levels. Consequently, this will also have effects on gene expression. However, the physical-chemical regulation of transcription is more complex than the initial idea that dense chromatin suppresses gene expression, while loose chromatin facilitates it^64,65^.

In summary, we have investigated and quantified the effect of ionic strengths, particularly divalent cations, and pH on the charge distribution of an NCP. We employed a molecular theory approach that was previously developed to study various interfacial polymer problems such as the ion-conductivity of polyelectrolytes of modified nanopores, the adsorption of proteins to polymer layers, and charge regulation of ligated nanoparticles^43,45,48^ The present paper extends and continues earlier studies on the charging behavior of biopolymer systems including nuclear core complexes, aggrecans, and bacteriophages, and as end-tethered polyelectrolytes.^43,46,47^. In this study, we explicitly account for the charge distribution of the AAs and DNA-phosphates of the nucleosome. Rather than making assumptions about the charging state of these components, our theory predicts their charged states based on their spatial distribution within the nucleosome.

Although we have added many molecular features, it should be realized that the theory is still an approximate mean-field approach that does not include electrostatic fluctuations. Another limitation of the theoretical approach is that we did not consider the dielectric property of the nucleosome. We assumed that the dielectric constant inside the nucleosome is equal to that of the background medium. Previous calculations for end-tethered polyelectrolyte layers, which used a local position-dependent dielectric constant (a linear volume-weighted average of the dielectric constants of water and the polymer, reflecting linear polarizability), found this to be a reasonable approximation for low and intermediate polymer densities^66^.

Recently, we applied a similar MT approach that includes a local position-dependent dielectric constant in combination with ion condensation to study the charging of self-assembled peptide amphiphiles (PA), which have a high density. The results indicated that using a local dielectric constant rather than a fixed one can lead to sizeable additional shifts in the charged state of the PA, particularly at pH values close to the pKa of the chargeable carboxylic acid. This effect was observed only in monovalent salt solutions. Since physiological pH values are outside the *pK*_*a*_ range of the AAs, and most charge regulation occurs via ion bridging in the lower-density regions of nucleosomes, assuming a fixed dielectric constant may be a reasonable initial approximation. Nonetheless, future studies should address the impact of the dielectric environment on the nucleosome’s charge state more thoroughly.

Another key assumption was that the distribution of AAs and DNA was considered rigid. Future research will relax this assumption and consider additional conformations of AA and DNA, especially that of the histone tails. Ion pairing reactions between AAs and phosphates were also not included, but since only about 1 in 10 phosphates are sufficiently close to interact with an AA, we believe this omission has minimal impact on the charging behavior. However, future work could explore this in greater detail, incorporating more molecular-level interactions into the theory to gain a better understanding of the charging characteristic of an NCP.

The single NCP model will be extended to include larger nucleosomal chains at higher concentrations, allowing us to model nucleosome interactions that are more representative of chromatin in both euchromatin and heterochromatin states. By increasing the complexity of the model to include chains of nucleosomes, we aim to capture variations in chromatin density and how these different configurations respond to environmental factors, such as ionic conditions and molecular crowding. This approach will enable us to investigate the contrasting behaviors between more loosely packed euchromatin and highly compacted heterochromatin. Specifically, we will explore how different levels of chromatin condensation in these two states affect the local intranuclear environment of nucleosomes. Through these extensions, we aim to better understand the structural implications in chromatin organization.

Additionally, although key ions like magnesium have been included in the theory, the nucleus contains a variety of other ions. In the future, multivalent ions such as polyamines will be added to create a more accurate representation of the nuclear environment. Given that the MT was originally developed to describe polymers^53^, this extension of the theory is feasible and actively being pursued. Thus, current work represents an initial step towards a more comprehensive understanding of charge regulation in chromatin, based on a detailed molecular description of the nucleosome’s constituent molecules.

## V. SUPPLEMENTARY MATERIAL

The supplementary material includes detailed derivations of the equations used in the Molecular Theory framework, additional results that complement the main findings, and specific computational details necessary for reproducing the analyses presented.

## ACKNOWLEDGMENTS

We acknowledge funding from the National Institutes of Health (NIH) grants U54CA268084, U54CA261694, 470 R01CA228272, R01CA224911, R01CA225002, T32GM142604, NSF grant EFMA-1830961, and philanthropic support from K. Hudson and R. Goldman, S. Brice and J. Esteve, M. E. Holliday and I. Schneider, the Christina Carinato Charitable Foundation and D. Sachs. Paola Carrillo Gonzalez acknowledges the support of NIH training grant T32GM142604. This research was supported in part through the computational resources and staff contributions provided for the Quest high performance computing facility at North-western University, which is jointly supported by the Office of the Provost, the Office for Research, and Northwestern University Information Technology.

The authors would like to thank Drs. Marcelo Carignano, Cody Dunton, and Luay Almassalha for useful discussions.

## AUTHOR DECLARATIONS

### Author Contributions

**Rikkert J. Nap**: Conceptualization (equal); Data curation (lead); Investigation (lead); Methodology (equal); Software (lead); Validation (equal); Visualization (equal); Writing – original draft (lead); Writing – review & editing (equal). **Paola Carillo Gonzalez**: Investigation (equal); Data curation (lead); Methodology (supporting); Software (equal); Validation (supporting); Visualization (equal); Writing – original draft (lead); Writing – review & editing (equal). **Aria Coraor**: Investigation (supporting); Methodology (supporting); Data curation (supporting); Software (supporting); Validation (supporting); Visualization (supporting);Writing – review & editing (equal) **Ranya K. A. Virk**: Conceptualization (equal); Investigation (support); Methodology (support); Software (support); Writing – review & editing (equal). **Juan de Pablo**: Conceptualization (supporting); Funding acquisition (supporting); Methodology (supoorting); Project administration (supporting); Resources (supporting) Supervision (lead); Writing – review & editing (equal). **Vadim Backman**: Conceptualization (equal); Funding acquisition (lead); Methodology (equal); Project administration (equal); Resources (lead); Supervision (lead); Writing – review & editing (equal). **Igal Szleifer**: Conceptualization (equal); Funding acquisition (lead); Methodology (equal); Project administration (lead); Resources (lead); Supervision (lead); Writing – review & editing (equal).

## DATA AVAILABILITY STATEMENT

The data that support the findings of this study are available from the corresponding author upon reasonable request.

