## Supplementary Material for "The Impact of Charge Regulation and Ionic Intranuclear Environment on the Nucleosome Core Particle"

[5)]*Chan Zuckerberg Biohub Chicago, Chicago, IL, USA*

[6)]*Department of Pharmaceutical Chemistry, University of California, San Francisco, CA, USA*

[7)]*Department of Chemistry, Northwestern University, Evanston, IL 60208, USA*[a)]

(*Electronic mail:)

[a)]† These authors contributed equally to this work.

### I. MOLECULAR THEORY

#### A. Divalent binding

A charged acid can bind with a divalent ion, like $Mg^{2+}$, in two distinct ways. Namely, one Mg-ion binds or condenses with one or two charged acids. The latter binding is referred to as bridging The following two reactions can represent this:

$$A^{-}(aq)+Mg^{2+}(aq) \rightleftarrows AMg^{+}(aq), \tag{S1}$$

$$A^{-}(aq)+AMg^{+}(aq)+ \rightleftarrows A_2Mg(aq). \tag{S2}$$

The latter reaction can also be represented thermodynamically equivalently like

$$2A^{-}(aq)+Mg^{2+}(aq)+ \rightleftarrows A_2Mg(aq). \tag{S3}$$

For an ideal solution, the equilibrium constant of this reaction is

$$K=\frac{[Mg^{2+}][A^{-}]}{[A_2Mg]}=\exp(-\beta\Delta G^{\ominus}). \tag{S4}$$

#### B. DNA-phosphate-Mg binding

First, focus on the metal-ion coordination or complexation of two charged phosphate acid molecules or monomers with one divalent Mg ion:

$$P_2Mg \rightleftarrows 2P^{-}+Mg^{2+}. \tag{S5}$$

Here, the objective here is to derive a free energy describing this multivalent ion binding process.

##### *1. Acid base equilibrium in end-tethered weak polyelectrolyte*

First, it is instructive to review the theory describing end-tethered weak polyelectrolytes. A detailed discussion of the theory can be found in Refs.[1,2]. The Helmholtz free energy of an end-tethered polyelectrolyte layer is given by

$$\beta F = \sum_{\alpha} P(\alpha) \ln P(\alpha) + \int d^3 r \rho_w(\vec{r}) \left(\ln(\rho_w(\vec{r}) v_w) - 1\right) \tag{S6}$$

$$+ \sum_{k \in \{K^+, Cl^-, OH^-, H^+\}} \int d^3 r \rho_k(\vec{r}) (\ln(\rho_k(\vec{r}) v_w) - 1 + \beta \mu_k^{\ominus}) \tag{S7}$$

$$+ \int d^3 r \langle \rho_p(\vec{r}) \rangle \left[ f_{AH}(\vec{r}) \left( \ln(f_{AH}(\vec{r}) + \beta \mu_{AH}^{\ominus} \right) + f_{A^-}(\vec{r}) \left( \ln(f_{A^-}(\vec{r}) + \beta \mu_{A^-}^{\ominus}) \right] \tag{S8}$$

$$+ \beta \int d^3 r \left( \langle \rho_q(\vec{r}) \rangle + \frac{1}{2} \varepsilon_0 \varepsilon_w (\nabla \psi(\vec{r}))^2 \right) \tag{S9}$$

here $\beta = 1/k_B T$ is the inverse temperature. The first term describes the conformational entropy of the end-tethered polymer layer. Here, $P(\alpha)$ is the probability of finding an end-tethered polymer chain in conformation $\alpha$. The thermodynamic and structural properties of the polymers can be calculated from the probability distribution function. For instance, the average number density of the polymer segments at position $\vec{r}$ is given by

$$\langle \rho_P(\vec{r}) \rangle = \sum_{\alpha} P(\alpha) n(\alpha; \vec{r}), \tag{S10}$$

here $n(\alpha; \vec{r}) d^3 r$ is the number of polymer segments within the element $[\vec{r}, \vec{r} + d\vec{r}]$ belonging to polymer conformation $\alpha$. Here, we assumed that the polyacid is a homopolymer.

The second term in the free energy expression represents the translational or mixing entropy of the solvent and $\rho_w(\vec{r})$ is the number density of the water (solvent) and $v_w$ is its volume. Then, the volume fraction of the water at $\vec{r}$ is $\phi_w(\vec{r}) = \rho_w(\vec{r}) v_w$. The next contributions in the free energy describe the mixing (translation) entropy and standard chemical potential of the mobile ionic species. Here, $\rho_i(\vec{r})$ is the number density of ionic species $i$ and the ion volume fraction is $\phi_i(\vec{r}) = \rho_i(\vec{r}) v_i$ with $v_i$ corresponding to its volume. In Eq. S6, $\mu_k^{\ominus}$ is the standard free energy of ionic species $k$.

The fourth term in the free energy describes the free energy associated with the chemical acid-base equilibrium

$$\mathrm{AH} \rightleftharpoons \mathrm{A^-} + \mathrm{H^+}, \tag{S11}$$

where $f_{AH}(\vec{r})$ corresponds to the fraction of protonated monomers at $\vec{r}$ and $f_{A^-}(\vec{r})$ is the fraction of monomers that is deprotonated/charged at $\vec{r}$. The first and third terms within the integral describe the entropy of the protonated charged state $AH$ and the deprotonated state ($A^-$) respectively. The second and fourth terms within the integral correspond to the standard chemical potential of the uncharged ($\mu_{AH}^{\ominus}$) and charged state ($\mu_{A^-}^{\ominus}$).

The fraction of protonated and deprotonated acid monomers add up to one at every position: $f_{A^-}(\vec{r}) + f_{AH}(\vec{r}) = 1$. Thus here $f_{AH}(\vec{r}) = 1 - f_{A^-}(\vec{r})$ and the separate introduction of the two fractions is not strictly necessary. However, it is convenient to introduce a fraction for every chemical state of the acid monomer when extended above free energy to include also monovalent sodium and potassium ion binding: $A^- + Na^+ \rightleftharpoons ANa$ and $A^- + K^+ \rightleftharpoons AK$. The inclusion of ion binding is then straightforward. For every chemical state one introduces an entropic and enthalpic free energy contribution, of the type similar to that of the (de) protonated states. These terms involve the fraction of the chemical states of the monomer. For example, the case of the potassium-acid ion state, it involves $f_{AK}(\vec{r})$.

The next two terms in free energy describe the electrostatic contribution to the free energy functional. Here $\psi(\vec{r})$ corresponds to the electrostatic potential at position $\vec{r}$ and $\langle\rho_q(\vec{r}\rangle$ is the charge density which is equal to

$$\langle\rho_q(\vec{r})\rangle = q_A f_{A^-}(\vec{r})\langle\rho_p(\vec{r})\rangle + \sum_{i=\{K,Cl,H,OH\}} q_i \rho_i(\vec{r}), \tag{S12}$$

where $q_i = z_i e$ is the elementary charge of ionic species $i$, with $z_i$ corresponding to its valence, and $q_A = -e$ the charge of the depronated acid.

Non-electrostatic van der Waals interactions ($E_{VdW}$ can be included in the free energy. Here for, simplicity, we can assume good solvent conditions; $E_{VdW} = 0$. The repulsive interactions in the theory are modeled as excluded volume interactions. The intrachain interactions are considered exactly during chain conformation generation, while intermolecular excluded volume interactions are accounted for by assuming that the system is incompressible at every position

$$\langle\phi_p(\vec{r}) + \phi_w(\vec{r}) + \sum_i \phi_i(\vec{r}) = 1. \tag{S13}$$

Here the first term corresponds to the total polymer volume fraction, which is given by

$$\langle\phi_p(\vec{r})\rangle = \langle\rho_p(\vec{r})\rangle \left(f_{A^-}(\vec{r})v_{A^-} + f_{AH}(\vec{r})v_{AH}\right). \tag{S14}$$

Here $v_{A^-}$ and $v_{AH}$ represented the volume of the acid monomer in the deprotonated and protonated chemical state respectively. The volume constraints are enforced by introducing Lagrange multipliers. Namely $\pi(\vec{r})$.

The free energy is minimized with respect to the $P(\alpha)$, $\rho_i(\vec{r})$, and $f_i(\vec{r})$, and varied with respect to the electrostatic potential $\psi(\vec{r})$ under the constraints of incompressibility, the fact that the system is in contact with a bath of cations, anions, protons, and hydroxyl ions and $\sum_i f_i(\vec{r}) = 1$.

Minimization of the free energy with respect to the solvent number density yields the following expression for the local volume fraction of the solvent

$$\phi_w(\vec{r}) = \rho_w(\vec{r})v_w = \exp\left(-\beta\pi(\vec{r})v_w\right), \tag{S15}$$

while minimization with respect to the ion density yield

$$\rho_\gamma(\vec{r}) = \frac{1}{v_w}\exp\left(\beta\mu_\gamma - \beta\mu_\gamma^{\ominus} - \beta\pi(\vec{r})v_\gamma - \beta\psi(\vec{r})z_\gamma e\right). \tag{S16}$$

Observe, the Lagrange multiplier, $\pi(\vec{r})$, can be interpreted as the lateral osmotic pressure. Also, the chemical potential of water is not specified explicitly because the incomprehensibility constraint reduces the number of independent thermodynamic variables. Therefore, the chemical potential, $\mu_\gamma$, is in reality an exchange chemical potential, i.e., the difference between the chemical potential of the molecule of type $\gamma$ and that of water. Likewise, the charge neutrality and the water self-dissociation equilibrium further reduce the number of independent thermodynamic variables. Values of the exchange chemical potentials of all species can be obtained by relating them to their reservoir concentrations: $\rho_\gamma^{bulk}v_w = \exp(\beta\mu_\gamma^{\ominus} - \beta\mu_\gamma - \beta\pi^{bulk}v_\gamma)$, with $\psi^{bulk} = 0$ (Eq. S71).[1,3].

Functional variation of the free energy with respect to the electrostatic potential yields a generalized Poisson equation and its boundary conditions. See Eq. S71.

Free energy minimizing results for the acid-base equilibrium into following reaction equations

$$\frac{f_{A^-}(\vec{r})}{f_{AH}(\vec{r})} = \frac{f_{A^-}(\vec{r})}{1-f_{A^+}(\vec{r})} = K_a^{\ominus}\frac{e^{-\beta\pi(\vec{r})\Delta v_a}}{\rho_{H+}(\vec{r})v_w}., \tag{S17}$$

Here $K_a^{\ominus} = \exp(-\beta\Delta G_a^{\ominus})$ c corresponds to the chemical equilibrium constant and $\Delta G_a^{\ominus})$ is the standard free energy change of either the acid-base equilibrium reaction of the acid, $\Delta v_A$ corresponds to the difference in volume between the products and the reactants. Explicitly, the standard free energy change of the acid-base equilibrium AH $\rightleftharpoons$ $A^- + H^+$<=> given by $\Delta G_a^{\ominus} = \mu_{A^-}^{\ominus} + \mu_{H^+}^{\ominus} - \mu_{AH}^{\ominus}$, and the change in volume is equal to $\Delta v_a = v_{A^-} + v_{H^+} - v_{AH}$ The chemical equilibrium constant $K_a^{\ominus}$ is related to the experimental acid–base equilibrium constant $K_a = C\exp(-beta\Delta D_a^{\ominus}$ of a single acidic monomer in infinitely dilute solution. Here $C$ is a constant required for the consistency of units and is equal to $C = 1/N_A v_w$, where $N_A$ is Avogadro's number.

Eq. S17 should be compared with the equation equilibrium constant of the single acid in aqueous solution.

$$K_a = \frac{[A^-][H^+]}{[AH]} \quad \text{equivalently} \quad \frac{f_{A^-}}{1-f_{A^-}} = \frac{K_a}{[H^+]}. \tag{S18}$$

The comparison shows that the chemical reaction equilibrium deviates from ideal charging directly via $e^{-\beta\pi(\vec{r})\Delta v_a}$ term in the numerator and position dependence of the local proton concentration, which obeys relation Eq. S16. The local proton concentration is influenced by excluded volume interactions as expressed by $\pi(\vec{r})$ and local electrostatic interactions as expressed by $\psi(\vec{r})$. Note that the electrostatic interaction is determined by the local charge density, which depends on the local chemical equilibrium. Thus, the local electrostatic interaction is coupled with the local chemical equilibrium as well as the excluded volume interactions. For an end-tethered polyacid layer this results in a significant reduction of charge state inside the polyacid layer. Further discussions of the theory and its implications for the charge and structural properties of polyacid layers can be found in reviews[1,2] and references therein.

Within the framework of molecular theory, we previously modeled divalent ion binding in context of end-tethered polyacids[4]. In that context, we introduce the fraction of acids that form a bridge involving two acids and one divalent ion, denoted as $f_{A_2Mg}(\vec{r})$. This fraction is analogous to those introduced earlier for the deprotonated, protonated, and monovalent ion condensed chemical states. Subsequently, we employed a mean-field approximation assuming either a lateral homogeneous distribution of the acids for the end-tethered polyacid layer with one or two spatial inhomogeneities, or a denser distribution of acids. This approximation did not differentiate between intra- versus inter-chain divalent ion bridges within an end-tethered polyacid layer, the latter were assumed to dominate. However, neither approximation is applicable to nucleosomes and NCPs. More specifically, the assumption of lateral homogeneity is absent for nucleosomes and NCPs. Unlike polyacids end-tethered to a surface, a nucleosome system lacks apparent symmetry and requires resolving the molecular theory in three spatial dimensions. Therefore, we developed a new and improved free energy description of ion-bridging applicable in 3D.

##### 2. *DNA-phosphate Pairs*

First, consider a solution of $N_p$ pairs of phosphate monomers. They are assumed to be immobile and have a fixed position. We assume that two phosphates that form a pair are sufficiently close that they can bridge and form an ion complex with $Mg^{2+}$.

The association free energy is then given as

$$F_{assoc} = N_p \left[ f_p(\ln f_p + \beta\mu^{\ominus}_{2P}) + (1-f_p)(\ln(1-f_p) + \beta\mu^{\ominus}_{PMg}) \right]. \tag{S19}$$

Here $f_p$ is the fraction of pairs not bound with a $Mg_2^+$ ion and $1 - f_p$ is the fraction of pairs bound with $Mg^{2+}$. Here $\mu^{\ominus}_{P_2Mg}$ corresponds to the standard free energy of the bound state, while $\mu^{\ominus}_{2P}$ is the standard free energy of the unbound pair of charged phosphates. The total free energy, assuming a fixed phosphate distribution, becomes

$$\begin{aligned}\frac{\beta W}{V} &= \sum_J \rho_J(\ln(\rho_J v_w) - 1 + \beta\mu_J^{\ominus}) + \rho_p\left[f_p(\ln(f_p) + \beta\mu^{\ominus}_{2P}) + (1-f_p)(\ln(1-f_p) + \beta\mu^{\ominus}_{P_2Mg})\right] \\ &+ \beta(\rho_q\psi - \frac{1}{2}\varepsilon_w\varepsilon_r(\nabla\psi)^2) + \beta\pi\left[\sum_J \rho_J v_J + \rho_p((1-f_p)v_{P_2Mg} + f_p)v_{2P}\right] \\ &-\beta\mu_{Mg^{2+}}(\rho_{Mg^{2+}} + \rho_p(1-f_p)) - \sum_{J\neq Mg^{2+}} \mu_J\rho_J \end{aligned} \tag{S20}$$

Here $\rho_{Mg^{2+}}$ corresponds to the density of free $Mg^{2+}$, while $\mu_{Mg^{2+}}(\rho_{Mg^{2+}} + \rho_p(1-f_p))$, occurring in last line of above the free energy, corresponds to the total number density of magnesium, which is the sum of free and bound magnesium. Minimization of the total free energy density with respect to $f_p$ under the constraint of mass balance yields the following reaction equation

$$\frac{f_p}{1-f_p} = K_d^{\ominus}\frac{e^{-\beta\pi\Delta v_d}}{\rho_{Mg^{2+}}v_w}, \tag{S21}$$

with

$$-\ln K_d^{\ominus} = \beta\Delta G_d^{\ominus} = \beta(\mu^{\ominus}_{2P-} + \mu^{\ominus}_{Mg^{2+}} - \mu^{\ominus}_{P_2Mg}), \tag{S22}$$

and

$$\Delta v_d = v_{2P-} + v_{Mg^{2+}} - v_{P_2Mg}. \tag{S23}$$

Here $v_{2P-}$ is the volume of one phosphate pair that has no bound magnesium, while $v_{P_2Mg}$ corresponds to the volume of a phosphate-magnesium bridge. Thus, $\Delta v_d$ corresponds to the change in volume as Mg binds the phosphate pair.

In Eq. S22, $\mu^{\ominus}_{2P-}$ is the standard free energy of one phosphate pair that has no bound magnesium, while $\mu^{\ominus}_{P_2Mg}$ corresponds to the standard free energy of a phosphate-magnesium bridge. Note for latter notation convenience we shall define $\mu^{\ominus}_{2P-} = 2\mu^{\ominus}_{P-}$. Here $\mu^{\ominus}_{P-}$ is half of the standard free energy of the phosphate pairs, for which both phosphates are charged. It is tempting to identify this with the standard free energy of a single charged phosphate implying that this free energy is twice the free energy of the two phosphates. This is not the case. The key approach here and in the next sections is that we treat the phosphate pairs as one, new particle or unit.

##### 3. *Pair density function*

Generalization by making $N_p$ position-dependent, i.e.:

$$N_p \quad \rightarrow \quad \langle \rho_p(\vec{r},\vec{r}')\rangle \mathrm{d}^3r\mathrm{d}^3r'. \tag{S24}$$

Here $\langle \rho_p(\vec{r},\vec{r}')\rangle \mathrm{d}^3r\mathrm{d}^3r'$ is the number of pairs with one element of the pair located at volume element $[\vec{r},\vec{r}+d\vec{r}]$ and the other element located within volume $[\vec{r}',\vec{r}'+d\vec{r}']$. This describes the pair density function of a phosphate pair located at $\vec{r}$ and $\vec{r}'$. Here $\langle A\rangle = \sum P(\alpha)A(\alpha)$ denotes thermodynamic averaging. Integration over both coordinates of the pair density function leads to the total number of pairs:

$$\frac{1}{2}\iint \mathrm{d}^3r\mathrm{d}^3r'\langle \rho_p(\vec{r},\vec{r}')\rangle = N_p. \tag{S25}$$

Here the half corrects for the doubling counting of the pairs. Here $f_p(\vec{r},\vec{r}')$, the generalization of $f_p$, is the fraction of pairs at $(\vec{r},\vec{r}')$ that are not bound with Mg. Then, the charge density becomes

$$\langle \rho_q(\vec{r})\rangle = \sum_i q\rho_i(\vec{r}) + \frac{1}{2}\int \mathrm{d}^3\mathrm{r}'\langle \rho_p(\vec{r},\vec{r}')\rangle f_p(\vec{r},\vec{r}')q_{P^-} + \frac{1}{2}\int \mathrm{d}^3\mathrm{r}'\langle \rho_p(\vec{r}',\vec{r})\rangle f_p(\vec{r}',\vec{r})q_{P^-}. \tag{S26}$$

Notice, that we do not consider the possibility of isolated phosphates: every phosphate has at least one phosphate with which it forms a pair. Using the definition of the pair density function, the phosphate volume fraction at position $\vec{r}$ equals

$$\begin{aligned}\langle \phi_P(\vec{r})\rangle = \frac{1}{2}\int \mathrm{d}^3\mathrm{r}'\langle \rho_p(\vec{r},\vec{r}')\rangle \left(f_p(\vec{r},\vec{r}')v_P + (1-f_p(\vec{r},\vec{r}'))v_{P_2Mg}/2\right) + \\ \frac{1}{2}\int \mathrm{d}^3\mathrm{r}'\langle \rho_p(\vec{r}',\vec{r})\rangle \left(f_p(\vec{r}',\vec{r})v_P + (1-f_p(\vec{r}',\vec{r}))v_{P_2Mg}/2\right).\end{aligned} \tag{S27}$$

The equation S19 for the chemical or association free energy becomes

$$\begin{aligned}F_{assoc} = \frac{1}{2}\iint \mathrm{d}^3r\mathrm{d}^3r'\langle \rho_p(\vec{r},\vec{r}')\rangle \big[f_p(\vec{r},\vec{r}')(\ln(f_p(\vec{r},\vec{r}')) + \beta\mu^{\ominus}_{2P}) \\ +(1-f_p(\vec{r},\vec{r}'))(\ln(1-f(\vec{r},\vec{r}')) + \beta\mu^{\ominus}_{P_2Mg})\big].\end{aligned} \tag{S28}$$

Minimization of the total free energy (see below) with respect to $f_p(\vec{r},\vec{r}')$ under the constraint of mass balance and incompressibility yield the following reaction equation

$$\frac{f_p(\vec{r},\vec{r}')}{1-f_p(\vec{r},\vec{r}')} = K_d^{\ominus}\frac{e^{-\frac{1}{2}\beta(\pi(\vec{r}')+\pi(\vec{r}))\Delta v_d}}{(\rho_{Mg^{2+}}(\vec{r})v_w)^{1/2}(\rho_{Mg^{2+}}(\vec{r}')v_w)^{1/2}}. \tag{S29}$$

Compare above equation with Eq. S21 and the equation previously obtained for the degree of deprotonation of acid monomers $AH \rightleftarrows A^- + H^+$

$$\frac{f_{A^-}(\vec{r})}{1-f_{A^-}(\vec{r})} = K_a^{\ominus}\frac{e^{-\beta\pi(\vec{r})\Delta v_a}}{\rho_{H^+}(\vec{r})v_w}. \tag{S30}$$

##### 4. *Free energy of nucleosome with only phosphate charge centers*

Here we first consider a nucleosome that has only DNA-phosphates charge centers. We consider this 'type' of macro-ion first to focus solely on the development and presentation of the Mg-pairing. In subsequent sections, we shall add the acid-base equilibrium and ion-pairing of the amino acids to the model. Here, the free energy pertains only to Mg-binding of a macro-ion or nucleosome that carries only DNA-phosphates charged. Here deprotonation and monovalent ion binding of the phosphates is not considered. The generalization and complete free energy of a nucleosome is presented in the next sections. Here in anticipation of the generalization and ease of mathematical derivation, we change the notation and denote the fraction of phosphate pairs that have (both) phosphates charged as $f_p(\vec{r},\vec{r}') \to f_{PP}(\vec{r},\vec{r}')$, and the fraction of phosphate pairs bound with Mg is denoted as $1-f_p(\vec{r},\vec{r}') \to f_{P_2Mg}(\vec{r},\vec{r}')$. The Lagrange multiplier $\lambda(\vec{r},\vec{r}')$ is introduced to enforce that $f_{PP}(\vec{r},\vec{r}')+f_{P_2Mg}(\vec{r},\vec{r}')=1$ for all $\vec{r}$ and $\vec{r}'$.

$$\begin{aligned}
\beta W &= \sum_\alpha P(\alpha)\ln P(\alpha) + \int \mathrm{d}^3r\rho_w(\vec{r})\left(\ln(\rho_w(\vec{r})v_w)-1\right) \\
&+ \sum_{k\in\{Na^+,K^+,Cl^-,OH^-,H^+,Mg^{2+}\}} \int d^3r\rho_k(\vec{r})(\ln(\rho_k(\vec{r})v_w)-1+\beta\mu_k^{\ominus}) \\
&+ \beta\int d^3r\left(\langle\rho_q(\vec{r})\rangle\psi(\vec{r})+\frac{1}{2}\varepsilon_0\varepsilon_w(\nabla\psi(\vec{r}))^2\right) \\
&+ \frac{1}{2}\iint \mathrm{d}^3r\mathrm{d}^3r'\langle\rho_p(\vec{r},\vec{r}')\rangle\left[f_{PP}(\vec{r},\vec{r}')(\ln(f_{PP}(\vec{r},\vec{r}'))+\beta\mu_{2P}^{\ominus})\right. \\
&\qquad \left.+f_{P_2Mg}(\vec{r},\vec{r}'))(\ln(f_{P_2Mg}(\vec{r},\vec{r}'))+\beta\mu_{P_2Mg}^{\ominus})\right]. \\
&+ \beta\frac{1}{2}\iint \mathrm{d}^3r\mathrm{d}^3r'\langle\rho_p(\vec{r},\vec{r}')\rangle\lambda(\vec{r},\vec{r}')\left[f_{PP}(\vec{r},\vec{r}')+f_{P_2Mg}(\vec{r},\vec{r}')-1\right] \\
&- \beta\mu_{Mg^{2+}}^{\ominus}\left(\int \mathrm{d}^3r\rho_{Mg^{2+}}(\vec{r})+\frac{1}{2}\iint \mathrm{d}^3r\mathrm{d}^3r'\langle\rho_p(\vec{r},\vec{r}')\rangle f_{P_2Mg}(\vec{r},\vec{r}')\right) \\
&- \sum_l\beta\mu_l\int \mathrm{d}^3r\rho_l(\vec{r})+\beta\int \mathrm{d}^3r\pi(\vec{r})\left(\langle\phi_P(\vec{r})\rangle+\sum_\gamma\rho_\gamma(\vec{r})v_\gamma-1\right)
\end{aligned} \tag{S31}$$

Minimization of $W$ with respect to $f_{PP}(\vec{r},\vec{r}')$ and $f_{P_2Mg}(\vec{r},\vec{r}')$ result in the following two equations

$$\beta\lambda(\vec{r},\vec{r}') = -\beta(\pi(\vec{r})+\pi(\vec{r}'))v_P-\beta(\psi(\vec{r})+\psi(\vec{r}'))q_P-\ln(f_{PP}(\vec{r},\vec{r}'))-2\beta\mu_{P^-}^{\ominus}-1 \tag{S32}$$

$$\beta\lambda(\vec{r},\vec{r}') = -\beta(\pi(\vec{r})+\pi(\vec{r}'))\frac{v_{P_2Mg}}{2}-\ln(f_{P_2Mg}(\vec{r},\vec{r}'))-\beta(\mu_{P_2Mg}^{\ominus}-\mu_{Mg^{2+}})-1 \tag{S33}$$

Combining the above equations and using the definitions for $\Delta G_d^{\ominus}$ and $\Delta v_d$ yield the chemical reaction equation of Eq. S29. Finally, the pair density is defined in terms of $P(\alpha)$ as follows

$$\langle \rho_p(\vec{r},\vec{r}') \rangle = \sum_\alpha P(\alpha)\rho_p(\alpha;\vec{r},\vec{r}'), \tag{S34}$$

here $\rho_p(\alpha;\vec{r},\vec{r}')\mathrm{d}^3r\mathrm{d}^3r'$ is the number of phosphate pairs located at $(\vec{r},\vec{r}')$ for a conformation $\alpha$. Since we consider only one nucleosome molecule, there are no pairs between different molecules. We only need to establish the number of intra-conformational pairs that exist in each conformation separately. The number of intra-conformational pairs can be established by counting the neighboring phosphates located within a certain distance that surrounds each phosphate:

$$\rho_p(\alpha;\vec{r},\vec{r}') \stackrel{def}{=} \frac{1}{2}\sum_{s\in S_{phosphate}} \left( \frac{n_s(\alpha;\vec{r},\vec{r}')}{\mathcal{N}_s(\alpha,\vec{r})} + \frac{n_s(\alpha;\vec{r}',\vec{r})}{\mathcal{N}_s(\alpha,\vec{r}')} \right), \tag{S35}$$

here $n_s(\alpha;\vec{r},\vec{r}')\mathrm{d}^3r\mathrm{d}^3r'$ is the number of neighboring phosphates that are located at $\vec{r}'$ relative to the phosphate of monomer number $s$ which is located at $\vec{r}(\alpha,s)$ that both belong to conformation $\alpha$. $S_{phosphate}$ denotes the set of monomer indices that label the DNA-phosphates. Mathematical $n_s(\alpha;\vec{r},\vec{r}')$ is given by

$$n_s(\alpha;\vec{r},\vec{r}') = \sum_{t\in S_{phosphate}, t\neq s} \delta^3(\vec{r}-\vec{r}(\alpha,s))\delta^3(\vec{r}'-\vec{r}(\alpha,t))U(\vec{r},\vec{r}'), \tag{S36}$$

with

$$U(\vec{r},\vec{r}') = \begin{cases} 1 & \text{if } |\vec{r}-\vec{r}'| \leq d_{cut} \\ 0 & \text{otherwise} \end{cases} \tag{S37}$$

Note there can be only one pair, hence we normalize $n_s(\alpha;\vec{r},\vec{r}')$ by the total number of neighbors in the volume shell $(\vec{r}')$, which is equal to $\mathcal{N}_s(\alpha,\vec{r}) = \int_{\partial V(\vec{r}=\vec{r}(\alpha,s)=\vec{r}'')}\mathrm{d}^3r'' \int \mathrm{d}^3r' n_s(\alpha;\vec{r}'',r')$. We only count those phosphates as a neighbor that are within a given cutoff distance from the central phosphate: $(|\vec{r}-\vec{r}'| \leq d_{cut})$. We choose $d_{cut} = 0.80\,nm$. This distance is motivated by the distance between two DNA-phosphate of neighboring base pairs along the same strand, which is roughly $0.72\,nm$ for DNA in a B-conformation. Larger distances are considered not to be able to form Mg-bridges. Consequently, pairing occurs primarily between nearest neighbor phosphates adjacent on the same strand of DNA. Bridging can also occur between non-neighbor phosphates. Since DNA is wrapped around the histone protein with around 1.5 turns. This brings several DNA-phosphates that are separated by one turn along the DNA chain to be within cutoff distance of each other.

Notice above equations, S35 and **??** count the number of neighboring phosphates for each phosphate in the conformation $\alpha$. This approximately equals the number of phosphate pairs of conformation. It constitutes a mean-field approximation, which is akin to the way we include acid-base equilibrium and ion-condensation in the MT approach.

In principle, the sum of conformations should include the sum for every spatial conformation ($\alpha$) and for each of the spatial conformation ($\alpha$) the number of unique ways of making Mg-bridges. The same applies to (de)protonation and ion binding. For every spatial state, we should count the number of chemical distinct states as well. Just considering the (de)protonation states would add $2^{N_D}$ additional conformations, with $N_{DNA}$ being the number of DNA-phosphates. This is clearly infeasible for any reasonable number of DNA-phosphates. Hence we employed the above approximation and accounted for the chemical equilibrium using the various fractions as detailed above. Further details on this aspect in the context of acid-base equilibrium of end-tethered polyacids can be found in e.g., Refs.[1–3].

Observe that, although this is a mean-field approximation the inclusion of conformations, i.e., spatial positions implies we have included spatial correlations, which go beyond traditional (monomeric) mean-field approaches. Similarly, by considering describing Mg-bridging through the "lens" of phosphate pairs, we also included the short-range correlations that are involved in Mg-bridging. In that sense, the developed scheme is reminiscent of a quasi-chemical approximation.

Finally, minimization with respect to $P(\alpha)$ yields

$$P(\alpha) = \frac{1}{q} \exp\left(\frac{1}{2}\beta \iint \mathrm{d}^3r \mathrm{d}^3r' \rho_p(\alpha;\vec{r},\vec{r}')\lambda(\vec{r},\vec{r}')\right), \tag{S38}$$

with $q$ ensuring that the pdf is properly normalized: $\sum_\alpha P(\alpha) = 1$.

#### *5. Generalization of Association Free energy*

Above free energy is restricted to the case that the phosphates of a pair are charged or form a $P_2Mg$ bridge. To account for protonation,

$$PH \rightleftarrows P^- + H^+, \tag{S39}$$

and monovalent ion binding,

$$PM \rightleftarrows P^- + M^+, \tag{S40}$$

we extend the above approach and introduce pair fractions for the case where the phosphates are both protonated $f_{(PH)(PH)}(\vec{r},\vec{r}')$ or one phosphate of the pairs is protonated $f_{(P)(PH)}(r,r'$ or

$f_{(PH)(P)}(\vec{r},\vec{r}')$ etc. Here, we use round brackets to emphasize the connection between phosphate type and its coordinate. Similarly, we introduce ion-binding pair fractions. Observed we can also have mixed pairs, for example, $f_{(PK)(PH)}(\vec{r},\vec{r}')$, which correspond to a phosphates pair where the phosphate located at $\vec{r}$ is bound with potassium and the phosphate at $\vec{r'}$. Generally, we denote these (monovalent) pair fractions as $f_{JK}(\vec{r},\vec{r}')$, with $J$ and $K$ denote the various (monovalent) chemical states of one of the phosphate of the phosphate ion pair. Namely

$$J,K \in \{P^-, PH, PNa, PK, PMg^+\}. \tag{S41}$$

Now the chemical free energy contribution reads

$$\beta F_{assoc} = \sum_{J,K} \frac{1}{2} \iint \mathrm{d}^3r \mathrm{d}^3r' \langle \rho_p(\vec{r},\vec{r}')\rangle \left\{ f_{JK}(\vec{r},\vec{r}') \left[ \ln(f_{JK}(\vec{r},\vec{r}')) + \beta(\mu_J^{\ominus} - \mu_K^{\ominus}) \right] \right. \tag{S42}$$

$$\left. + f_{P_2Mg}(\vec{r},\vec{r}') \left( \ln(f_{P_2Mg}(\vec{r},\vec{r}')) + \beta \mu^{\ominus}_{P_2Mg} \right) \right\}, \tag{S43}$$

and the charge density becomes

$$\langle \rho_q(\vec{r})\rangle = \sum_i q_i \rho_i(\vec{r}) + \frac{1}{2} \int \mathrm{d}^3\mathrm{r}' \langle \rho_p(\vec{r},\vec{r}')\rangle \sum_{J,K} f_{JK}(\vec{r},\vec{r}') q_J + \frac{1}{2} \int \mathrm{d}^3\mathrm{r}' \langle \rho_p(\vec{r}',\vec{r})\rangle \sum_{J,K} f_{JK}(\vec{r}',\vec{r}) q_K, \tag{S44}$$

with $\{q_{P^-}, q_{PH}, q_{PNa}, q_{PK}, q_{PMg^+}\} = \{-e, 0, 0, 0, +e\}$, and the phosphate volume fraction becomes

$$\langle \phi_{phos}(\vec{r})\rangle = \frac{1}{2} \int \mathrm{d}^3\mathrm{r}' \langle \rho_p(\vec{r},\vec{r}')\rangle \left( \sum_{J,K} f_{JK}(\vec{r},\vec{r}') v_J + f_{P_2Mg}(\vec{r},\vec{r}') v_{P_2Mg}/2 \right)$$
$$+ \frac{1}{2} \int \mathrm{d}^3\mathrm{r}' \langle \rho_p(\vec{r}',\vec{r})\rangle \left( \sum_{JK} f_{JK}(\vec{r}',\vec{r}) v_K + f_{P_2Mg}(\vec{r}',\vec{r}) v_{P_2Mg}/2 \right). \tag{S45}$$

##### *6. Chemical Equilibrium Equations*

The total free energy see below, is minimized with respect to all the different monovalent pair fractions, $f_{JK}(\vec{r},\vec{r}')$, and the fraction of the Mg-bridges, $f_{P_2Mg}(\vec{r},\vec{r}')$. Minimization with respect of $f_{JK}(\vec{r},\vec{r}')$, meaning:

$$\frac{\delta W}{\delta f_{JK}(\vec{r},\vec{r}')} = 0, \tag{S46}$$

yields

$$\ln(f_{JK}(\vec{r},\vec{r}')) + \beta(\mu_J^{\ominus} + \mu_K^{\ominus}) + \beta(\pi(\vec{r}) v_J + \pi(\vec{r}')) v_K + \beta(\psi(\vec{r}) q_J + \psi(\vec{r}') q_K) +$$
$$- \sum_{M \in \{H^+, Na^+, K^+, Mg^2+\}} \beta \mu_M (\delta_J^{PM} + \delta_K^{PM}) + 1 + \beta \lambda(\vec{r},\vec{r}') = 0. \tag{S47}$$

We can now readily solve for $f_{JK}(\vec{r},\vec{r}'))$ by eliminating $\lambda(\vec{r},\vec{r}')$ by subtracting above equation from the equation for the case $JK = (P)(P) = PP$. This is similar to the procedure used in section I B 4. The result for $JK = (PH)(PH)$ is

$$\frac{f_{PP}(\vec{r},\vec{r}')}{f_{(PH)(PH)}(\vec{r},\vec{r}')} = K_a^{\ominus} \frac{e^{-\beta\pi(\vec{r})\Delta v_a}}{\rho_{H^+}(\vec{r})v_w} \cdot K_a^{\ominus} \frac{e^{-\beta\pi(r')\Delta v_a}}{\rho_{H^+}(\vec{r}')v_w}, \tag{S48}$$

the case $JK = (PH)(P)$ gives

$$\frac{f_{PP}(\vec{r},\vec{r}')}{f_{(PH)(P)}(\vec{r},\vec{r}')} = K_a^{\ominus} \frac{e^{-\beta\pi(\vec{r})\Delta v_a}}{\rho_{H^+}(\vec{r})v_w}, \tag{S49}$$

whereas $JK = (P)(PH)$ gives

$$\frac{f_{PP}(\vec{r},\vec{r}')}{f_{(P)(PH)}(\vec{r},\vec{r}')} = K_a^{\ominus} \frac{e^{-\beta\pi(\vec{r}')\Delta v_a}}{\rho_{H^+}(\vec{r}')v_w}. \tag{S50}$$

Here $K_a^{\ominus}$ is the equilibrium constant of the deprotonation of one of the phosphate of the phosphate pair:

$$-\ln K_a^{\ominus} = \beta\Delta G_a^{\ominus} = \beta(\mu_{P^-}^{\ominus} + \mu_{H^+}^{\ominus} - \mu_{PH}^{\ominus}), \tag{S51}$$

and $\Delta v_a$ is the change in volume upon deprotonation of one of the phosphate of the phosphate pair

$$\Delta v_a = v_{P^-} + v_{H^+} - v_{PH}. \tag{S52}$$

The general case of $JK = (PM)(PL)$ yields

$$\frac{f_{PP}(\vec{r},\vec{r}')}{f_{(PM)(PL)}(\vec{r},\vec{r}')} = K_m^{\ominus} \frac{e^{-\beta\pi(\vec{r})\Delta v_m}}{\rho_{M^+}(\vec{r})v_w} \cdot K_l^{\ominus} \frac{e^{-\beta\pi(\vec{r}')\Delta v_l}}{\rho_{L^+}(\vec{r}')v_w}. \tag{S53}$$

Here $K_m^{\ominus}$ and $\Delta v_m$ are defined in completely analogous way to $K_a^{\ominus}$ and $\Delta v_a$. The equation of the Mg-bridge is

$$\frac{f_{PP}(\vec{r},\vec{r}')}{f_{P_2Mg}(\vec{r},\vec{r}')} = K_d^{\ominus} \frac{e^{-\frac{1}{2}\beta(\pi(\vec{r}')+\pi(\vec{r}))\Delta v_d}}{(\rho_{Mg^{2+}}(\vec{r})v_w)^{1/2}(\rho_{Mg^{2+}}(\vec{r}')v_w)^{1/2}}. \tag{S54}$$

Equations S48-S54 correspond to the ratio between the number of pairs in chemical state $(P)(P)$, i.e., both phosphates charged and the number of pairs in chemical state $(PM)(PL)$ or the Mg-bridge, chemical state $(P_2Mg)$, respectively. With these ratios, we can solve for $f_{JK}(\vec{r},\vec{r}')$ and $f_{P_2Mg}(\vec{r},\vec{r}')$ by noting that the sum over all different chemical states of the pairs needs to add up to one for every position: $\sum_{JK} f_{JK}(\vec{r},\vec{r}') + f_{P_2Mg}(\vec{r},\vec{r}') = 1$. It is convenient to define the

$$x_{JK}(\vec{r},\vec{r}') \stackrel{def}{=} \frac{f_{JK}(\vec{r},\vec{r}')}{f_{PP}(\vec{r},\vec{r}')} \tag{S55}$$

This is the inverse of ratios in Eqs. S48-S54, Now $f_{PP}$ becomes

$$f_{PP}(\vec{r},\vec{r}') = \frac{1}{1+\sum^{*}_{J,K} x_{JK}(\vec{r},\vec{r}') + x_{P2Mg}(\vec{r},\vec{r}')} \tag{S56}$$

The star in the summation signifies that the summation exclude the state $JK = (P)(P)$. The remaining chemical fractions are given by

$$f_{JK}(\vec{r},\vec{r}') = x_{JK}(\vec{r},\vec{r}') f_{PP}(\vec{r},\vec{r}'). \tag{S57}$$

Observe that the ratios are dependent on $\pi(\vec{r})$ and $\psi(\vec{r})$ only. Hence the fractions are determined uniquely in terms of $\pi(\vec{r})$ and $\psi(\vec{r})$. The final expression for the ratios presented in S48-S54 resulted from applying the definitions of $\Delta K_k^{\ominus}$ and rewriting the terms involving the lateral pressure and electrostatic potential in terms for the ion density of $Na^+$, $K^+$, and $Mg^{2+}$.

$$\rho_k(\vec{r}) = \frac{1}{v_w} exp(-\beta(\mu_k^{\ominus} - \mu_k) - \pi(\vec{r})v_k - \psi(\vec{r})q_k) \tag{S58}$$

This leads to the expression presented in Eqs. S48-S54.

#### C. Free energy of a single NCP

Combining the association free energy for DNA-phosphates with the free energy with association free energies that involves the acid-base equilibrium and ion binding of the amino acid results in the following Helmholtz free energy for a single NCP

$$\begin{aligned}
\beta F &= \sum_{\alpha} P(\alpha)\ln P(\alpha) + \int d^3r \rho_w(\vec{r}) \left(\ln(\rho_w(\vec{r})v_w) - 1\right) \\
&+ \sum_{k\in\{Na^+,K^+,Cl^-,OH^-,H^+,Mg^+\}} \int d^3r \rho_k(\vec{r})(\ln(\rho_k(\vec{r})v_w) - 1 + \beta\mu_k^{\ominus}) \\
&+ \sum_{A\in\{AAacids\}} \int d^3r \langle\rho_A^C(\vec{r})\rangle \left[f_{AH}(\vec{r})\left(\ln(f_{AH}(\vec{r}) + \beta\mu_{AH}^{\ominus}\right) + f_{A^-}(\vec{r})\left(\ln(f_{A^-}(\vec{r}) + \beta\mu_{A^-}^{\ominus}\right)\right. \\
&+ \left. f_{ANa}(\vec{r})\left(\ln(f_{ANa}(\vec{r}) + \beta\mu_{ANa}^{\ominus}\right) + f_{AK}(\vec{r})\left(\ln(f_{AK}(\vec{r}) + \beta\mu_{AK}^{\ominus}\right)\right] \\
&+ \sum_{B\in\{AAbases\}} \int d^3r \langle\rho_B^C(\vec{r})\rangle \left[f_{BH^+}(\vec{r})\left(\ln(f_{BH^+}(\vec{r}) + \beta\mu_{BH^+}^{\ominus}\right) + f_B(\vec{r})\left(\ln(f_B(\vec{r}) + \beta\mu_B^{\ominus}\right)\right. \\
&+ \left. f_{BHCl}(\vec{r})\left(\ln(f_{BHCl}(\vec{r}) + \beta\mu_{BHCl}^{\ominus}\right)\right] \\
&+ \sum_{J,K}\frac{1}{2}\iint \mathrm{d}^3r\mathrm{d}^3r' \langle\rho_p(\vec{r},\vec{r}')\rangle \left\{f_{JK}(\vec{r},\vec{r}')\left[\ln(f_{JK}(\vec{r},\vec{r}')) + \beta(\mu_J^{\ominus} - \mu_K^{\ominus})\right]\right. \\
&\qquad + \left. f_{P_2Mg}(\vec{r},\vec{r}')\left(\ln(f_{P_2Mg}(\vec{r},\vec{r}')) + \beta\mu_{P_2Mg}^{\ominus}\right)\right\} \\
&+ \beta \int d^3r \left(\langle\rho_q(\vec{r})\rangle + \frac{1}{2}\varepsilon_0\varepsilon_w(\nabla\psi(\vec{r}))^2\right).
\end{aligned} \tag{S59}$$

where, $\langle\rho_q(\vec{r})\rangle$ corresponds to the total charge number density, which is equal to

$$\langle\rho_q(\vec{r})\rangle = \sum_k q_k\rho_k(\vec{r}) + \sum_{A\in\{Asp,Cys,Glu,Tyr\}} \langle\rho_A^C(\vec{r})\rangle f_{A^-}(\vec{r})(-e) + \sum_{B\in\{Arg,His,Lys\}} \langle\rho_B^C(\vec{r})\rangle f_{BH^+}(\vec{r})(+e)$$
$$+\frac{1}{2}\int \mathrm{d}^3\mathrm{r}'\langle\rho_p(r,r')\rangle \sum_{J,K} f_{JK}(\vec{r},\vec{r}')q_J + \frac{1}{2}\int \mathrm{d}^3\mathrm{r}'\langle\rho_p(\vec{r}',\vec{r})\rangle \sum_{J,K} f_{JK}(\vec{r}',\vec{r})q_K. \quad \text{(S60)}$$

Here, in the free energy and the charge density, $\langle\rho_A^C(\vec{r})\rangle$ denotes the number density of the chargeable acid centers of an acidic amino acids, with A corresponding to Asp, Cys, Glu, or the Tyr amino acid. Similar, $\langle\rho_B^C(\vec{r})\rangle$ corresponds to the number density of the chargeable centers of the basic amino acids. Namely Arg, His, and Lys. $f_{A^-}(\vec{r})$ is the fraction of acidic AA residues that are charged or deprotonated at position $\vec{r}$, while $f_{AH}(\vec{r})$ is the fraction of neutral, protonated acids, and $f_{ANa}(\vec{r})$ and $f_{AK}(\vec{r})$ are the fraction of acids that are condensed with $Na^+$ or $K^+$, respectively. Similarly, $f_{BH^+}(\vec{r})$ is the fraction of basic amino-acids residues that are charged or protonated at position $r$, $f_B(\vec{r})$ is the fraction of neutral, deprotonated basic amino acid bases, and $f_{BHCl}(\vec{r})$ is the fraction of basic amino acids that are condensed with $Cl^-$.

The total free energy is minimized with respect to the number density of all species, $\rho_k(\vec{r})$, the fraction of the charged states, $f_k(\vec{r})$, and varied with respect to the electrostatic potential, $\psi(\vec{r})$, under the constraints of incompressibility and the fact that the system is in contact with a bath of cations, anions, protons, and hydroxide ions. Thus, the proper thermodynamic potential is the semi-grand potential: $W = F - \sum_i \mu_i N_i$, with $N_i$ denoting the total number of particles of species $i$.

$$\begin{aligned}
\beta W &= \beta F + \beta\int \mathrm{d}^3 r\pi(\vec{r})\left(\langle\phi_{DNA}(\vec{r})\rangle + \sum_A\langle\phi_A(\vec{r})\rangle + \sum_B\langle\phi_B(\vec{r})\rangle + \sum_N\langle\phi_N(\vec{r})\rangle + \sum_k \rho_k(\vec{r})v_k - 1\right) \\
&+ \sum_{M\in\{H^+,Na^+,K^+,Mg^2+\}} -\beta\mu_M\left(\int \mathrm{d}^3 r\rho_M(\vec{r}) + \sum_{J,K}\frac{1}{2}\iint \mathrm{d}^3 r\mathrm{d}^3 r'\langle\rho_p(\vec{r},\vec{r}')\rangle f_{JK}(\vec{r},\vec{r}')\left(\delta_J^{PM}+\delta_K^{PM}\right)\right. \\
&+ \left.\sum_{A\in\{AA,acids\}}\int d^3 r\langle\rho_A^C(\vec{r})\rangle f_{AM}(\vec{r}) + \delta_M^{Mg^{2+}}\frac{1}{2}\iint \mathrm{d}^3 r\mathrm{d}^3 r'\langle\rho_p(\vec{r},\vec{r}')\rangle f_{P_2Mg}(\vec{r},\vec{r}')\right) \\
&- \beta\mu_{OH^-}\int \mathrm{d}^3 r\rho_{OH^-}(\vec{r}) - \beta\mu_{Cl^-}\left(\int \mathrm{d}^3 r\rho_{Cl^-}(\vec{r}) + \sum_{B\in\{AA,bases\}}\int d^3 r\langle\rho_B^C(\vec{r})\rangle f_{BHCl}(\vec{r})\right) \\
&+ \beta\frac{1}{2}\iint \mathrm{d}^3 r\mathrm{d}^3 r'\langle\rho_p(\vec{r},\vec{r}')\rangle\lambda(\vec{r},\vec{r}')\left[\sum_{J,K} f_{JK}(\vec{r},\vec{r}') + f_{P_2Mg}(\vec{r},\vec{r}') - 1\right] \\
&+ \beta\int \mathrm{d}^3\langle\rho_A^C(\vec{r})\rangle\lambda_A(\vec{r})\left[f_{A^-}(\vec{r}) + f_{AH}(\vec{r}) + f_{AK}(\vec{r}) + f_{ANa}(\vec{r}) - 1\right] \\
&+ \beta\int \mathrm{d}^3\langle\rho_B^C(\vec{r})\rangle\lambda_B(\vec{r})\left[f_{BH^+}(\vec{r}) + f_B(\vec{r}) + f_{BHCl}(\vec{r}) - 1\right]
\end{aligned} \quad \text{(S61)}$$

Here, $\pi(\vec{r})$ are Lagrange multipliers that enforce the packing constraint, while the flowing three lines ensure mass conservation. The last three lines ensure that the sum over all chemical states of the phosphate pairs, acidic and basic amino acids at every position add up to one. Explicitly the incompressibility constraint reads:

$$\langle \phi_{nuc}(\vec{r}) \rangle + \phi_w(\vec{r}) + \sum_{\gamma} \rho_{\gamma}(\vec{r}) v_{\gamma} = 1, \tag{S62}$$

The first term corresponds to the total volume fraction of the nucleosome, the second is the volume fraction of water, and the last term corresponds to the sum of volume fractions of all ionic mobile species.

##### *1. Number density and volume fraction and molecular representation NCP*

The total volume fraction of the nucleosome is the sum of the contributions arising from the DNA, the acidic AAs, basic AAs, and neutral AAs and is given by

$$\langle \phi_{nuc}(\vec{r}) \rangle = \langle \phi_{DNA}(\vec{r}) \rangle + \sum_{A} \langle \phi_A(\vec{r}) \rangle + \sum_{B} \langle \phi_B(\vec{r}) + \sum_{N} \langle \phi_N(\vec{r}) \rangle. \tag{S63}$$

As outlined in the main text and the section "Chain Generation", within the model, DNA is atomistically represented by a triplet of units, namely a chargeable phosphate (P), a sugar (S), and a nucleobase, either (C, T, A, or G) using the 3SPN.2 model developed by the de Pablo group[5]. The histone core (octamer) proteins, composed of AAs, are represented at the atomic level. Each atom in the side chain of an AA is depicted as a single unit. Assuming the histone protein is in a frozen 'crystal' state we can use the atom positions determined from X-ray diffraction experiments directly in our model[6].

Using this representation of the NCP/nucleosome the volume fraction of DNA becomes

$$\langle \phi_{DNA}(\vec{r}) \rangle = \sum_{s=\{C,T,A,G,S\}} \langle \rho_{DNA,s}(\vec{r}) \rangle v_s^{DNA} + \langle \phi_{phos}(\vec{r}) \rangle, \tag{S64}$$

with $\langle \phi_{phos}(\vec{r}) \rangle$ is the volume fraction of the phosphates as given by Eq. 21 in the main text and Eq. S45. Here $\rho_{DNA,s}(\vec{r}) \rangle$ and $v_s^{DNA}$ correspond to the number density and volume of either sugar or nucleobase. Observe that depending on the charged state of the phosphate pairs the total volume fraction of the phosphates and DNA changes as can be seen for Eq. $S45$. Similarly, the volume fraction of neutral amino acids

$$\langle \phi_N(\vec{r}) \rangle = \sum_{s} \langle \rho_{N,s}(\vec{r}) \rangle v_{N,s}, \tag{S65}$$

where the index $s$ runs over all atom units of the AAs. The volume fraction of the acidic AA is defined as

$$\langle\phi_A(\vec{r})\rangle = \sum_s \langle\rho_{A,s}(\vec{r})\rangle v_{A,s} + \langle\rho_A^C(\vec{r})\rangle \left(f_{A^-}(\vec{r})v_{A^-} + f_{AH}(\vec{r})v_{AH} + f_{AK}(\vec{r})v_{AK} + f_{ANa}(\vec{r})v_{ANa}\right). \tag{S66}$$

Here the index $s$ runs over all neutral atom units of the AAs. Similarly, the volume fraction of the basic AAs is defined ad

$$\langle\phi_B(\vec{r})\rangle = \sum_k \langle\rho_{B,s}(\vec{r})\rangle v_{B,s} + \langle\rho_B^C(\vec{r})\rangle \left(f_{BH^+}(\vec{r})v_{BH^+} + f_B(\vec{r})v_B + f_{BHCl}(\vec{r})v_{BHCl}\right). \tag{S67}$$

Considering multiple different NCP conformations, would in analog to previous MT approaches, yield for the number densities of the various densities of DNA and AAs.

$$\langle\rho_{\gamma,s}(\vec{r})\rangle = \sum_\alpha P(\alpha) n_{\gamma,s}(\alpha;\vec{r}). \tag{S68}$$

Here $n_{\gamma,s}(\alpha;\vec{r})d^3r$ is the number of 'atom' units of type $(\gamma,s)$ located within volume element $[\vec{r},\vec{r}+d\vec{r}]$ belonging to conformation $\alpha$. Compare this to the definition of the pair density of phosphates ( Eq. S34) and the definition of the polymer density of an end-tethered polyacid layer (Eq. S10).

Next, we assume that the NCP is in one 'frozen' state represented by one conformation only. Therefore in the remainder of the text, we will omit the conformational dependency and write in general $A$ instead of $\langle A\rangle$. Thus, for example $n_{\gamma,s}(\alpha;\vec{r})$ becomes $n_{\gamma,s}(\vec{r})$. The same applies to the pair density of Eq. S34.

#### 2. *Free energy minimization*

Minimization of the free energy with respect to the solvent number density yields the following expression for the local volume fraction of the solvent

$$\phi_w(\vec{r}) = \rho_w(\vec{r})v_w = \exp\left(-\beta\pi(\vec{r})v_w\right), \tag{S69}$$

while minimization with respect to the ion density yield

$$\rho_\gamma(\vec{r}) = \frac{1}{v_w}\exp\left(\beta\mu_\gamma - \beta\mu_\gamma^\ominus - \beta\pi(\vec{r})v_\gamma - \beta\psi(\vec{r})z_\gamma e\right). \tag{S70}$$

This equation is identical to the equation presented previously in Eqs. S15 and S16.

Functional variation of the free energy with respect to the electrostatic potential yields a generalized Poisson equation and its boundary conditions

$$-\varepsilon_0 \varepsilon_w \nabla^2 \psi(\vec{r}) = \rho_q(\vec{r}) \quad \nabla \cdot \psi(\vec{r})|_{R\to\infty} = 0 \quad \psi(R \to \infty) = \psi^{bulk} = 0 \tag{S71}$$

Minimization of the free energy with respect to $f_{JK}$ and $f_{P2Mg}$, the different chemical states of the phosphate pair yield the equations presented in the foregoing section. Minimization of the free energy with respect to the fraction of (de)protonated and ion-condensed states of the acidic amino acids yields

$$\frac{f_{A^-}(\vec{r})}{f_{AH}(\vec{r})} = K_a^{\ominus} \frac{e^{-\beta(\pi(\vec{r})\Delta v_a}}{\rho_{H+}(\vec{r}) v_w}, \tag{S72}$$

and

$$\frac{f_{A^-}(\vec{r})}{f_{AM}(\vec{r})} = K_d^{\ominus}(M) \frac{e^{-\beta(\pi(\vec{r})\Delta v_d}}{\rho_{M^+}(\vec{r}) v_w}. \tag{S73}$$

Similarly, minimization of the free energy with respect to the fraction of (de)protonated and ion-condensed states of the basic amino acids yields

$$\frac{f_B(\vec{r})}{f_{BH^+}(\vec{r})} = K_a^{\ominus} \frac{e^{-\beta(\pi(\vec{r})\Delta v_a}}{\rho_{H+}(\vec{r}) v_w}, \tag{S74}$$

and

$$\frac{f_{BH^+}(\vec{r})}{f_{BHCl}(\vec{r})} = K_d^{\ominus}(Cl) \frac{e^{-\beta(\pi(\vec{r})\Delta v_{BHCl}}}{\rho_{Cl^-}(\vec{r}) v_w}. \tag{S75}$$

Note, like the fractions for the different chemical states of the phosphate pair, we can solve readily for the fraction of different chemical states and express them as a function of $\pi(\vec{i})$ and $\psi(\vec{i})$. Defining $x_{AH}(\vec{r}) = \frac{f_{AH}(\vec{r}}{f_{A^-}(\vec{r})}$ and $x_{AM}(\vec{r}) = \frac{f_{AM}(\vec{r})}{f_{A^-}(\vec{r})}$ and noting that the sum over all different chemical states of the acidic AA adds up to one for every position: $\sum_{A\in\{A^-,AH,AK,ANa\}} f_A(\vec{r}) = 1$ gives

$$f_{A^-}(\vec{r}) = \frac{1}{1 + x_{AH}(\vec{r}) + x_{AK}(\vec{r}) + x_{ANa}(\vec{r})} \tag{S76}$$

The remaining fractions are given by $f_{AH}(\vec{r}) = f_{A^-}(\vec{r}) x_{AH}(\vec{r})$, $f_{AK}(\vec{r}) = f_{A^-}(\vec{r}) x_{AK}(\vec{r})$, and $f_{ANa}(\vec{r}) = f_{A^-}(\vec{r}) x_{ANa}(\vec{r})$. Similar expressions hold for the basic amino acids.

#### D. Chain Generation

The result presented in the main text pertains only to one nuclear core particle. Here, for completeness and with the future extension towards oligonucleosomes in mind, we outline the general procedure to generate a representative set of (unbiased) conformations representing an oligonucleosome, i.e., multiple nucleosomes, using coarse-grained molecular dynamics simulations.

Oligonucleosome conformations for Molecular Theory calculations were produced by coarse-grained molecular dynamics in LAMMPS[7] using a modification of the 1-Cylinder-per-Nucleosome (1CPN) model[8]. This model represents each nucleosome as a rigid, aspherical particle with an anisotropic interaction potential, which captures the free energy of nucleosome-nucleosome interactions, and represents linker DNA as a discrete, charged, twistable, and stretchable wormlike chain at a 3-base pair-per-bead level of coarse-graining[8]. After trajectory generation, we map the 1CPN representation of the nucleosome to that of 3SPN for DNA and a fully atomistic representation for the histone octamers employed by MT.

To create appropriately unbiased starting structures for analysis in Molecular Theory, electrostatics in these simulations were removed by replacing the repulsive Coulomb interaction between DNA beads with an exclusively repulsive excluded volume (Lennard-Jones) term, and the magnitude of attractive internucleosome interactions were scaled down to $0.05kcal/mol$ to enforce excluded volume without including significant electrostatic-based attraction. Other parameters were left unchanged.

An octamer of nucleosomes connected with 50-base pair linkers (i.e., NRL 197) was generated by the initialization scripts provided by 1CPN. Unbiased coarse-grained MD simulations were performed until a total of N conformations, across 5 independent replicas, were obtained, for a total simulation time of $750\mu s$. Simulations were performed without periodic boundaries at a constant temperature of 300K enforced with a Langevin thermostat. The resulting conformations were then converted from the 1CPN representation to the 3SPN/AICG representation using a custom Python script (made available on GitHub). For use in the Molecular Theory calculations, the AICG ($\alpha$-carbon) representation was converted into a 'fully' atomistic representation with regard to the histone octamer by using the relative atom positions of the 'heavy' atoms, whose positions were determined using X-ray diffraction experimental data[6]. Finally, we used experimental X-ray observations by[9,10] to identify which part of the histone tails are in a disordered state. Those amino acid residues were removed from the representation of the no-tail configurations.

Finally, to remove the sampling bias effect caused by the small remaining attractive internucleosomal interactions, and obtain an accurate, unbiased distribution of the conformational space, we reweighted each conformation based on the energy derived from the nonbonded potential. The weight of a particular conformation in relation to the set of all conformations is $\mathrm{w_{conf}} = \frac{e^{E_{conf}}}{\sum_i e^{E_i}}$.

This appropriately reweights the trajectory's ensembles to the thermodynamic ensemble of the same system simulated without the attractive potential. This reweighting procedure is expected to have minimal effect on final observations.

##### E. Numerical methodology

Substituting the equations for the NCP, solvent, and ion volume fractions. Eqs. S63-S67, S69, and S70, into the packing constraint, Eq. S62, the generalized Poisson equation, Eq. S71, combined with the chemical reaction equations, Eqs. S72-S75 and Eqs. S48-S50 and Eqs. S53-S54, results in a set of coupled integro-differential equations. The unknowns of these equations are the position-dependent lateral pressure $\pi(\vec{r})$ and electrostatic potential $\psi(\vec{r})$. A numerical solution for the lateral pressure and electrostatic potential is obtained by discretization of the packing constraints, Eq. S62, and the generalized Poisson equation, Eq. S71. The equations are discretized by dividing the $\vec{r}$-coordinates into cubic cells of dimension $\delta$. Position-dependent functions are assumed to be constant within a cell, hence integrations can be replaced by summations. The integral of a general position-dependent function $f(\vec{r})$ then becomes:

$$\int_V \mathrm{d}^3 r f(\vec{r}) = \sum_{i,j,k} \int_{(i-1)\delta}^{i\delta} \mathrm{d}x \int_{(j-1)\delta}^{j\delta} \mathrm{d}x \int_{(k-1)\delta}^{k\delta} \mathrm{d}z\, f(x,y,z) \approx \delta^3 \sum_{i,j,k} f(i,j,k) \tag{S77}$$

Here $f(i,j,k)$ denotes the value which function $f(\vec{r}) = f(x,y,z)$ attains within the cubic region located between $(i-1)\delta \leq x < i\delta$, $(j-1)\delta \leq y < j\delta$, and $(k-1)\delta \leq z < k\delta$.

The packing constraint, Eq. S62, in discrete form for grid cell $(i,j,k)$ reads

$$\phi_{nuc}(i,j,k)\rangle + \phi_w(i,j,k) + \phi_{K^+}(i,j,k) + \phi_{Cl^-}(i,j,k) \\ + \phi_{Na^+}(i,j,k) + \phi_{H^+}(i,j,k) + \phi_{OH^-}(i,j,k) + \phi_{Mg^{2+}}(i,j,k) = 1. \tag{S78}$$

The volume fraction of water, Eq. S69, in discrete form becomes

$$\phi_w(i,j,k) = \exp(-\beta\pi(i,j,k)v_w), \tag{S79}$$

while the volume fraction of the counter ions, co ions, protons, and hydroxyl ions, Eq. S70, in discretized space are:

$$\phi_{Na^+}(i,j,k) = \phi_{Na^+,bulk}\exp(-\beta(\pi(i,j,k)-\pi_{bulk})v_{Na^+} - e\beta\psi(i,j,k)), \quad (S80)$$

$$\phi_{K^+}(i,j,k) = \phi_{K^+,bulk}\exp(-\beta(\pi(i,j,k)-\pi_{bulk})v_{K^+} - e\beta\psi(i,j,k)), \quad (S81)$$

$$\phi_{Cl^-}(i,j,k) = \phi_{Cl^-,bulk}\exp(-\beta(\pi(i,j,k)-\pi_{bulk})v_{Cl^-} + e\beta\psi(i,j,k)), \quad (S82)$$

$$\phi_{H^+}(i,j,k) = \phi_{H^+,bulk}\exp(-\beta(\pi(i,j,k)-\pi_{bulk})v_{H^+} - e\beta\psi(i,j,k)), \quad (S83)$$

$$\phi_{OH^-}(i,j,k) = \phi_{OH^-,bulk}\exp(-\beta(\pi(i,j,k)-\pi_{bulk})v_{OH^-} + e\beta\psi(i,j,k)). \quad (S84)$$

$$\phi_{Mg^{2+}}(i,jk) = \phi_{Ca^{2+},bulk}\exp(-\beta(\pi(i,j,k)-\pi_{bulk})v_{Mg^{2+}} - 2e\beta\psi(i,j,k)). \quad (S85)$$

The above volume fractions depend on the lateral pressure, electrostatic potential, and bulk volume fractions of the mobile ions. The chemical potentials of the counter and co-ions, protons, and hydroxyl ions are related to their bulk volume fractions.[3] These bulk values are input to the theory.

The volume fraction of the NPC and the related densities of AAs and phosphates in discrete form are :

$$\phi_{nuc}(i,j,k) = \phi_{DNA}(i,j,k) + \sum_A \phi_A(i,j,k) + \sum_B \phi_B(i,j,k) + \sum_N \phi_N(i,j,k), \quad (S86)$$

with

$$\phi_{DNA}(i,j,k) = \sum_{k=\{C,T,A,G,S\}} \rho_{DNA,s}(i,j,k)v_s^{DNA} + \phi_{phos}(i,j,k). \quad (S87)$$

The volume fraction of the neutral AAs is

$$\phi_N(i,j,k) = \sum_s \rho_{N,s}(i,j,k)v_{N,s}. \quad (S88)$$

The volume fraction of the acidic AAs in discrete form is

$$\phi_A(i,j,k) = \sum_s \rho_{A,s}(i,j,k)v_{A,s} + \rho_A^C(i,j,k)\left(f_{A^-}(i,j,k)v_{A^-} + f_{AH}(i,j,k)v_{AH} + f_{AK}(i,j,k)v_{AK} + f_{ANa}(i,j,k)v_{ANa}\right). \quad (S89)$$

Similarly, the volume fraction of the basic AAs in discrete form is

$$\phi_B(i,j,k) = \sum_s \rho_{B,s}(i,j,k)v_{B,s} + \rho_B^C(i,j,k)\left(f_{BH^+}(i,j,k)v_{BH^+} + f_B(i,j,kv_B + f_{BHCl}(i,j,k)v_{BHCl}\right). \quad (S90)$$

The densities occurring in the expressions for the DNA and AA volume fractions, Eqs. S87-S90, in discrete form are

$$\rho_{\gamma,s}(i,j,k) = \frac{1}{\delta^3}\int_{(i-1)\delta}^{i\delta}\mathrm{d}x\int_{(j-1)\delta}^{j\delta}\mathrm{d}y\int_{(k-1)\delta}^{k\delta}\mathrm{d}z\rho_{\gamma,s}(x,y,z) \tag{S91}$$

In a way similar to the definition of AA densities, the phosphates pair density, $\rho_p(\vec{r},\vec{r}')$ (Eq. S92), in discrete form is

$$\begin{aligned}\rho_p(\vec{i},\vec{i}') \stackrel{def}{=} \rho_p(i,j,k;i',j',k') = \frac{1}{\delta^6}&\int_{(i-1)\delta}^{i\delta}\mathrm{d}x\int_{(j-1)\delta}^{j\delta}\mathrm{d}y\int_{(k-1)\delta}^{k\delta}\mathrm{d}z \\ &\times\int_{(i'-1)\delta}^{i'\delta}\mathrm{d}x'\int_{(j'-1)\delta}^{j'\delta}\mathrm{d}y'\int_{(k'-1)\delta}^{k'\delta}\mathrm{d}z'\rho_p(x,y,z,x',y',z').\end{aligned} \tag{S92}$$

Here, we introduced the notation $\vec{i}$ to denote the triplet of indexes $(i,j,k)$. With the above definitions, the phosphate volume fraction, Eq. S45 becomes

$$\begin{aligned}\phi_{phos}(\vec{i}) = &\frac{\delta^3}{2}\sum_{\vec{j}}\rho_p(\vec{i},\vec{j})\left(\sum_{J,K} f_{JK}(\vec{i},\vec{j})v_J + f_{P_2Mg}(\vec{i},\vec{j})v_{P_2Mg}/2\right) \\ &+\frac{\delta^3}{2}\sum_{\vec{j}}\rho_p(\vec{j},\vec{i})\left(\sum_{JK} f_{JK}(\vec{j},\vec{i})v_K + f_{P_2Mg}(\vec{j},\vec{i})v_{P_2Mg}/2\right).\end{aligned} \tag{S93}$$

and the phosphate charge density becomes

$$\rho_{q,phos}(\vec{i}) = \frac{\delta^3}{2}\sum_{\vec{j}}\rho_p(\vec{i},\vec{j})\sum_{J,K} f_{JK}(\vec{i},\vec{j})q_J + \frac{\delta^3}{2}\sum_{\vec{j}}\rho_p(\vec{j},\vec{i})\sum_{J,K} f_{JK}(\vec{j},\vec{i})q_K. \tag{S94}$$

The chemical reaction equations related to the different chemical states of the phosphate pairs are listed below. The equation of the fraction of Mg-bridges is

$$\frac{f_{PP}(\vec{i},\vec{j})}{f_{P_2Mg}(\vec{i},\vec{j})} = K_d^{\ominus}\frac{e^{-\frac{1}{2}\beta(\pi(\vec{i})+\pi(\vec{j}))\Delta v_d}}{(\rho_{Mg^{2+}}(\vec{i})v_w)^{1/2}(\rho_{Mg^{2+}}(\vec{j}')v_w)^{1/2}}. \tag{S95}$$

The equations for the fraction of phosphate pairs in chemical state $JK = (PM)(PL)$ is

$$\frac{f_{PP}(\vec{i},\vec{j})}{f_{(PM)(PL)}(\vec{i},\vec{j})} = K_m^{\ominus}\frac{e^{-\beta\pi(\vec{i})\Delta v_m}}{\rho_{M^+}(\vec{i})v_w}\cdot K_l^{\ominus}\frac{e^{-\beta\pi(\vec{j})\Delta v_l}}{\rho_{L^+}(\vec{j})v_w}. \tag{S96}$$

The equation for the fraction of phosphate pairs in either a protonated or partial protonated chemical state, $(PH)(P)$ or $(P)(PH)$, can be readily obtained from Eq. S96.

The chemical reaction equations related to the acidic AAs, Eqs. S72 and S73, are in discrete form

$$\frac{f_{A^-}(i,j,k)}{f_{AH}(i,j,k)} = K^{\ominus}_{AH}\frac{e^{-\beta\pi(i,j,k)\Delta v_{AH}}}{\rho_{H^+}(i,j,k)v_w}, \quad \text{(S97)}$$

$$\frac{f_{A^-}(i,j,k)}{f_{ANa}(i,j,k)} = K^{\ominus}_{ANa}\frac{e^{-\beta\pi(i,j,k)\Delta v_{ANa}}}{\rho_{Na^+}(i,j,k)v_w}, \quad \text{(S98)}$$

$$\frac{f_{A^-}(i,j,k)}{f_{AK}(i,j,k)} = K^{\ominus}_{AK}\frac{e^{-\beta\pi(i,j,k)\Delta v_{AK}}}{\rho_{K^+}(i,j,k)v_w}. \quad \text{(S99)}$$

Similarly, the chemical reaction equations, related to the basic AAs, Eqs. S74, and S75, are in discrete form

$$\frac{f_B(i,j,k)}{f_{BH^+}(i,j,k)} = K^{\ominus}_{BH^+}\frac{e^{-\beta(\pi(i,j,k)\Delta v_{BH^+}}}{\rho_{H^+}(i,j,k)v_w}, \quad \text{(S100)}$$

and

$$\frac{f_{BH^+}(i,j,k)}{f_{BHCl}(i,j,k)} = K^{\ominus}_{d}(Cl)\frac{e^{-\beta(\pi(i,j,k)\Delta v_{BHCl}}}{\rho_{Cl^-}(i,j,k)v_w}. \quad \text{(S101)}$$

Using the finite volume method of box integration method, the Poisson equation, in cartesian coordinates, takes on the following discretized form

$$\psi(i+1,j,k)+\psi(i-1,j,k)+\psi(i,j+1,k)+\psi(i,j-1,k)+\psi(i,j,k+1)+\psi(i,j,k-1) - 6\psi(i,j,k) = -\varepsilon_w\varepsilon_0\delta^3\rho_q(i,j,k), \quad \text{(S102)}$$

here the total charged density is equal to

$$\rho_q(\vec{i}) = \sum_k q_k\rho_k(\vec{i}) + \sum_{A\in\{Asp,Cys,Glu,Tyr\}} \rho^C_A(\vec{r})f_{A^-}(\vec{i})(-e) + \sum_{B\in\{Arg,His,Lys\}} \rho^C_B(\vec{i})f_{BH^+}(\vec{i})(+e) + \rho_{q,phos}(\vec{i}). \quad \text{(S103)}$$

The discretization scheme applies only to the interior grid cells. In a similar way, we can construct, using the finite volume method, discretization schemes for the Poisson equation for the boundaries, i.e., the edges and corners of the grid. For example the discretization of the Poisson equation at the edge, $x=0$, of the grid becomes using the electrostatic boundary condition for the surface $x=0$, yields

$$\psi(2,j,k)+\psi(1,j+1,k)+\psi(1,j-1,k)+\psi(1,j,k+1)+\psi(1,j,k-1) - 5\psi(1,j,k) = -\varepsilon_w\varepsilon_0\delta^3\rho_q(1,j,k). \quad \text{(S104)}$$

Further details on numerical discretization schemes can be found in Refs.[11,12].

The discretization of the packing constraints and generalized Poisson equation turn these integro-differential equations into a set of coupled nonlinear algebraic equations that can be solved by standard numerical methods for example[13].

#### F. Composition of nucleosome

TABLE S1. Number of chargeable aminio acids and phosphate of (1kx5) nucleosome with and with out tails. The last column corresponds to the $pK_a$. The first five moieties are acids, while the last three are bases.

| | no tails | with tails | $pK_a$ |
|---|---|---|---|
| Asp | 22 | 24 | 3.71 |
| Cys | 2 | 2 | 8.14 |
| Glu | 48 | 48 | 4.15 |
| P | 292 | 292 | 1.0 |
| Tyr | 30 | 30 | 10.1 |
| Arg | 94 | 106 | 12.1 |
| His | 18 | 18 | 6.04 |
| Lys | 80 | 114 | 10.67 |

TABLE S2. Volume of nucleosome components. The volumes were determined based on the minimum atomic distances to ensure no overlap between atoms, thus avoiding steric hindrances.

| Atom Type | Volume ($nm^3$) |
|---|---|
| Phosphate | 0.025 |
| Sugar | 0.025 |
| Adenine | 0.021 |
| Cytosine | 0.024 |
| Guanine | 0.021 |
| Thymine | 0.024 |
| AA side-chain | 0.010 |

TABLE S3. Volume of solvent and ions in the system. Volumes were determined based on ionic radii. For ion condensation reactions, the volume of the bound product is the sum of volumes ($v_{AM} = v_A + v_M$). For acid-base reactions the volume of the protonated and deprotonated state is the same ($v_{AH} = v_{A^-}$ and $v_{BH^+} = v_B$).

| | Volume ($nm^3$) |
|---|---|
| $K^+$ | 0.011 |
| $Na^+$ | 0.0045 |
| $Mg^{2+}$ | 0.0016 |
| $Cl^-$ | 0.025 |
| w | 0.03 |
| $H^+$ | 0.03 |
| $OH^-$ | 0.03 |

#### G. Additional Results

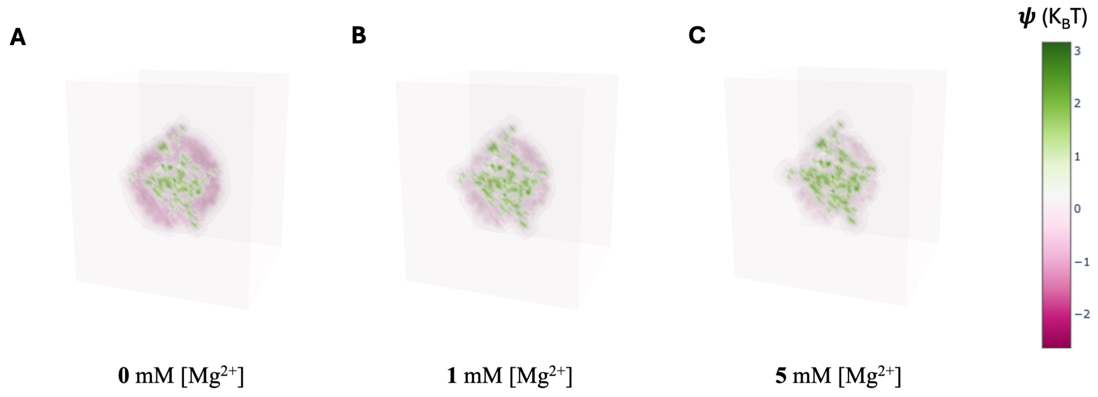


FIG. S1. 3D electrostatic potential plots of a nucleosome lacking disordered histone tails at varying $Mg^{2+}$ concentrations. As $Mg^{2+}$ concentration increases, the DNA (depicted in pink with negative electrostatic potential) becomes progressively neutralized. Salt concentrations of the other ions in the system are as follows: [KCl]=140 mM, and [NaCl]=10 mM

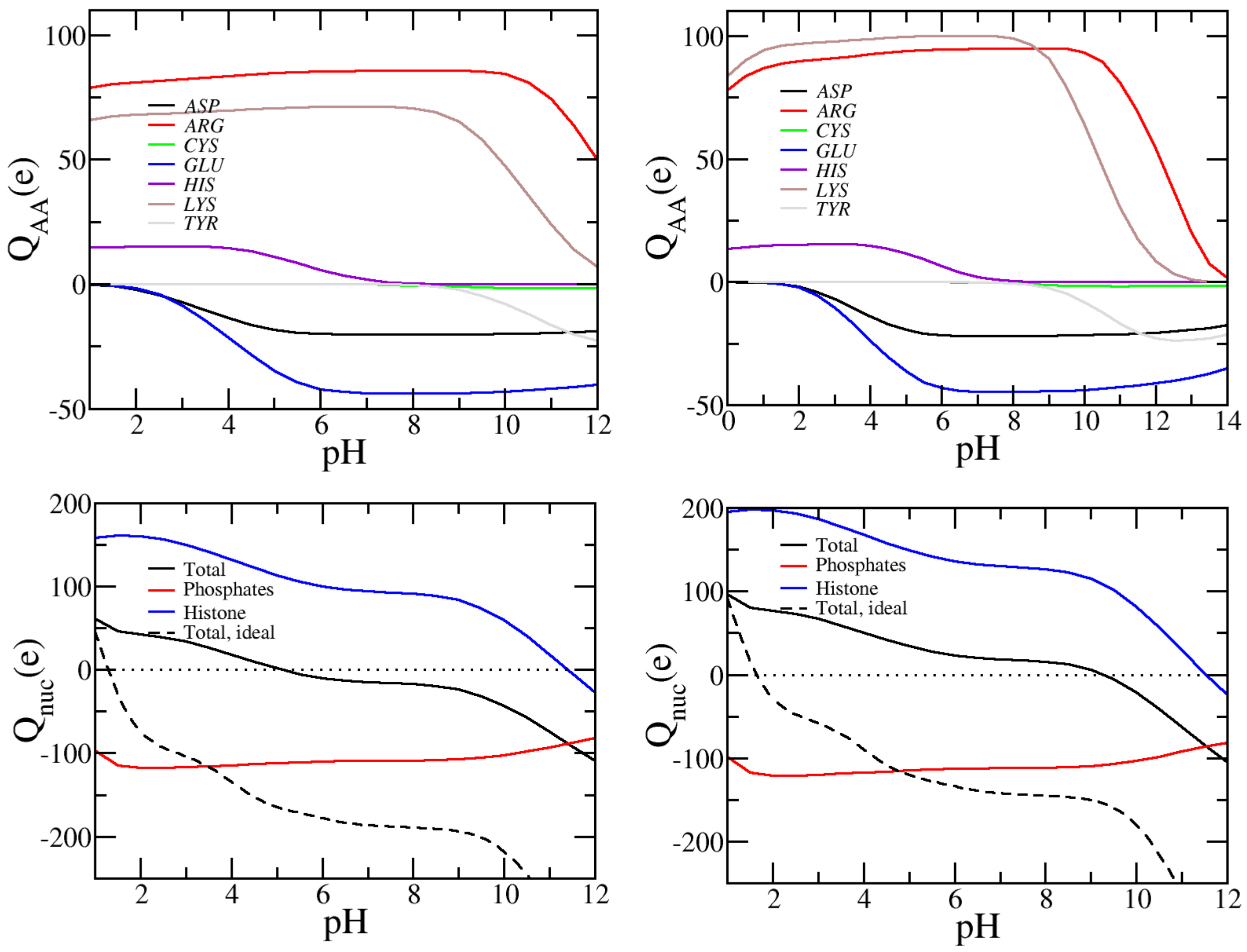


FIG. S2. The top row shows the charge contribution of amino acids in the system with regards to pH for a nucleosome with (right) and without (left) disordered histone tails. The bottom row shows the charge contribution of the phosphates, the total nucleosome charge, and net amino acids charge with regards to pH for a nucleosome with (right) and without (left) disordered histone tails. The dashed line shows the total ideal charge of the nucleosome. The ideal charge only considers the acid-base equilibrium of the amino acids and phosphates using the Henderson-Hasselbalch equation. It does not account for monovalent ion condensation or Mg-bridging. Comparison of MT predicted charge and ideal charge emphasizes again the large effect of Mg and ion condensation on the net charge of a nucleosome. Salt concentrations are as follows: [KCl]=140 mM, [NaCl]=10 mM, and [$MgCl_2$]=1 mM.

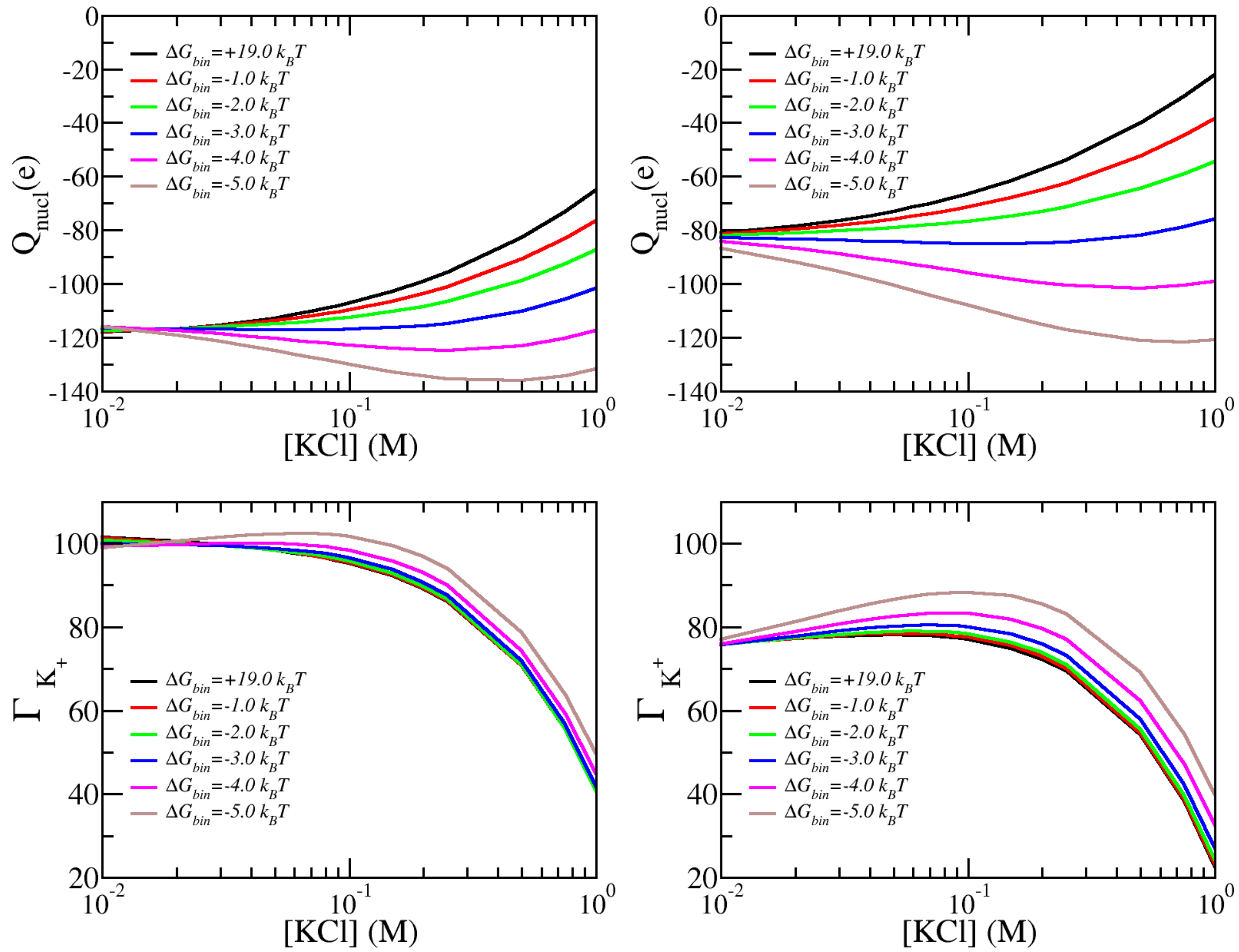


FIG. S3. The total effective charge, $Q_{nucl}$ for a nucleosome without and with tails as function of KCl concentration for physiological condition $pH = 7.4$ and [$MgCl_2$]=0 M.

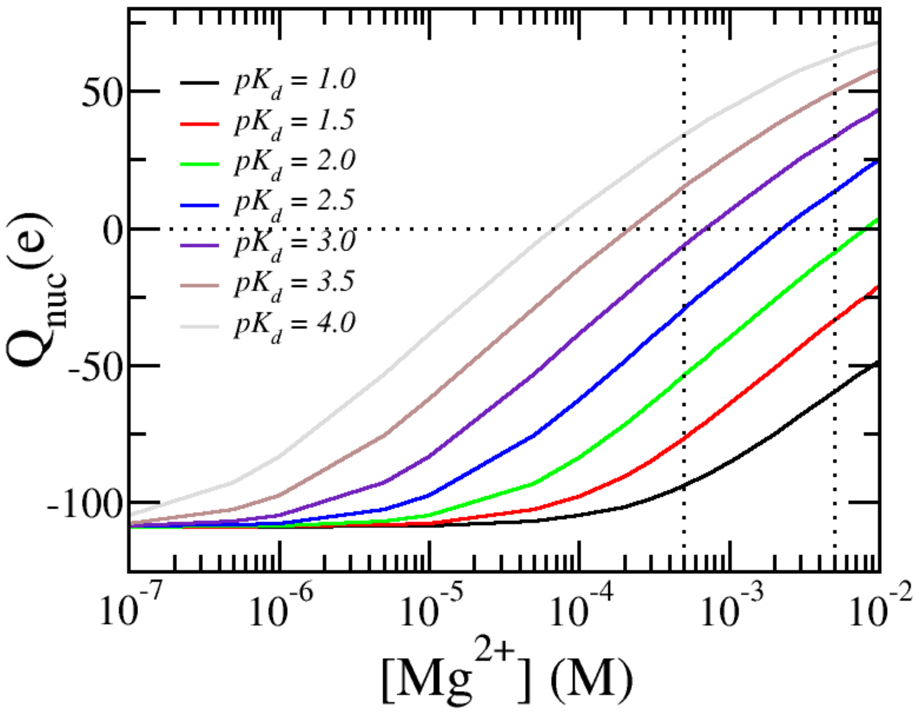


FIG. S4. The total effective charge for a nucleosome without histone tails considering different $P2Mg$ dissociation constants ($pK_d$).
